# Heart-On-a-Chip with Integrated Ultrasoft Mechanosensors for Continuous Measurement of Cell- and Tissue-scale Contractile Stresses

**DOI:** 10.1101/2025.10.16.682609

**Authors:** Ali Mousavi, Christina-Marie Boghdady, Shihao Cui, Sabra Rostami, Amid Shakeri, Naimeh Rafatian, Mark Aurousseau, Gregor Andelfinger, Milica Radisic, Christopher Moraes, Houman Savoji

**Author notes:** **Corresponding Author: Houman Savoji** - Institute of Biomedical Engineering, Department of Pharmacology and Physiology, Faculty of Medicine, University of Montréal, Montréal, QC, Canada; CHU Sainte-Justine Research Center, Montréal, QC, Canada; Montréal TransMedTech Institute (iTMT), Montréal, QC, Canada.

## Abstract

Heart-on-a-chip platforms aim to miniaturize and replicate the complex structure and function of cardiac tissue. Traditionally, microfabricated pillar pairs have been employed in these systems to provide tissue anchorage and determine contractility parameters based on pillar deflection. However, this approach lacks the spatial resolution required to capture local cell- and tissue-scale mechanical stresses. In this study, we established a non-destructive optical method for continuous micro- and macro-scale contractile force measurements. We utilized our previously developed edge-labeled micro-spherical stress gauges (eMSGs) to map the stresses within a heart-on-a-chip. These ultrasoft mechanosensors visibly deform in response to stresses generated by cells and the extracellular matrix (ECM). The chip consisted of two cell-seeding chambers, each containing flexible silicone pillar pairs to support tissue formation and compaction. Neonatal rat cardiomyocytes (CMs) were encapsulated in a fibrin/Geltrex hydrogel mixture containing eMSGs and seeded into each chamber. Over time, the tissue compacted and began beating spontaneously, demonstrating structural alignment and functional cardiac hallmarks, such as calcium transients and tissue-scale beating. The effects of ECM composition on tissue function were examined, revealing that lower fibrin concentrations significantly enhanced contractile frequency, regularity, and stress generation. Local cell- and ECM-scale mechanics were further investigated by analyzing the shape changes of the dispersible sensors. Lateral and longitudinal stresses were calculated for each sensor, highlighting the critical role of tissue compaction and contraction in cell-generated forces. Finally, the platform was validated using two known drug candidates, with their effects on contractility clearly demonstrated.

## 1. Introduction

Cardiovascular disease (CVD) is the leading cause of death globally [1]. Contractile dysfunction is a key hallmark of many CVDs, primarily leading to a reduced pumping capacity of the heart and, ultimately, heart failure.[2]. Since cardiomyocytes (CMs) are the contractile cells of the heart muscle (i.e., myocardium) and are primarily responsible for the heart’s functionality, investigating their contractility *in vitro* is crucial. Such studies are essential for developing drugs and therapeutic methods and advancing our understanding of the cellular and molecular mechanisms underlying CVDs. [3]. The heart-on-a-chip (HOC) platform has been introduced as a versatile three-dimensional (3D) *in vitro* model designed to replicate the complex network of the native heart, including cellular distribution, tissue dynamics, and key extracellular matrix (ECM) properties such as structural, compositional, electrophysiological, and biomechanical cues [4]. This biomimetic device can generate physiologically relevant tissue functionality, such as calcium transients and synchronous beating, making it a valuable tool for examining and validating drug safety and efficacy during the preclinical stage of the drug development pipeline [5].

Various methods have been reported for measuring CM contractility *in vitro*, including traction force microscopy (TFM) [6], atomic force microscopy (AFM) [7], optical imaging (e.g., video-based detection of cell edge displacement or sarcomere shortening) [8], electrical impedance spectroscopy [9], and the deflection of well-defined anchoring structures such as posts [10], cantilevers [11], or wires [12, 13]. Traditional methods, such as AFM and TFM, require specialized and expensive equipment and have primarily been used for single-cell analysis or two-dimensional (2D) cell sheets. This limitation restricts their applicability for measuring contractility in 3D tissues [14]. As an alternative model, engineered heart tissues (EHTs) have been developed by incorporating CMs into a hydrogel matrix, which gradually compacts over time into specific shapes, such as tissue strips or rings [15]. Tissue compaction requires the presence of stromal cells, such as fibroblasts. The resulting tissue shape is influenced by the mold design, the number of anchoring points, and the seeding pattern. The hydrogels most commonly used as ECM components for EHTs include collagen, Matrigel, fibrin, or blends of these materials [16].

While rigid anchoring structures, such as metal or glass rods and stainless-steel needles, were used in early EHT studies, the use of flexible pillars made from silicon-based elastomers, such as polydimethylsiloxane (PDMS), offers significant advantages. These flexible pillars allow for continuous measurement of tissue contractile activity by analyzing their deflection patterns and mechanical properties [17]. However, these quantified forces only reflect global tissue-scale contractility and fail to capture the local heterogeneity of cell-generated forces.

Cell-generated forces play a pivotal role in shaping tissue structure, determining cell fate, and regulating functionality during various stages of tissue remodeling and morphogenesis. These forces are crucial from the developmental stage through the maintenance of homeostasis and into the progression of diseases [18]. The critical role of these forces has been extensively studied in various biological and biomedical processes, including cell migration [19], matrix compaction [20], muscle contraction [21], wound healing [22], and cancer progression [23, 24]. Despite their importance, measuring cell-scale forces and monitoring force variations at local supracellular scales within dynamic 3D live tissues remain significant challenging. Addressing this issue requires a non-destructive, real-time measurement method. Dispersible, cell-sized mechanosensors have emerged as a promising solution to overcome these challenges [25, 26].

Biopolymer-based sensors can be fabricated as either oil droplets or hydrogel droplets and can be embedded into 3D engineered tissues *in vitro* or animal models *in vivo*. These sensors are capable of measuring various anisotropic or isotropic internal stresses, including compression and tension, depending on their specific design and intended application [27, 28]. We and others [25, 29, 30] have recently introduced ultrasoft, biocompatible, and fluorescently-labelled hydrogel droplets with well-defined mechanical properties. When integrated into a 3D matrix, imaging the shapes of these sensors allows us to infer local cell-generated stresses via computational analysis. In previous work, the Moraes group modified this approach to create edge-labelled microspherical stress gauges (eMSGs) that allows for long-term, stable fluorescent imaging, and used these to study engineered tumor models *in vitro* and *in vivo* [24].

Most previous HOC systems focused exclusively on global contractility and did not explore cell-scale tissue mechanics. In this study, we addressed this limitation by combining the aforementioned force measurement techniques through the integration of eMSGs into a functional HOC model. The chip containsflexible pillar pairs, enabling the assessment of tissue-scale contractile forces through pillar deflection. Simultaneously, the deformation of eMSGs allowed real-time measurement of local cell-scale stresses, providing a comprehensive view of contractile dynamics across scales (i.e., 10s of microns to mm).

## 2. Materials and Methods

### 2.1. Materials

All chemical reagents were supplied by Sigma Aldrich, Canada, unless otherwise specified. Polyglycerol polyricinoleate (PGPR 4150) was purchased from Palsgaard, Denmark. Carboxylate-modified polystyrene nanoparticles were obtained from ThermoFisher, Canada. Bovine collagen type I was supplied by Advanced Biomatrix, USA. Sulfo-SANPAH was purchased from G-Biosciences, USA. Mathematical modeling and computations were carried out using COMSOL Multiphysics v6.2 (COMSOL Inc., USA), and MATLAB R2023b (The Mathworks Inc., USA).

#### 2.1.1. Materials for Cell-based Experiments

Dulbecco’s modified Eagle medium (DMEM)/F12 culture medium (#10565018), Dulbecco’s phosphate buffered saline (DPBS; without calcium and magnesium), fetal bovine serum (FBS, #16140089), penicillin/streptomycin (Pen Strep, #15140122), Geltrex basement membrane matrix (#A1413201), trypan blue (#15250061), GlutaMAX (#35050061) were purchased from Gibco, Canada. Claycomb medium (#51800C), fibrinogen (#F8630), thrombin (#T4648), aprotinin (#A3428), Pluronic-F127, bovine serum albumin (BSA), paraformaldehyde solution (PFA, 4% in PBS), norepinephrine (NE), blebbistatin (Bleb), and dimethyl sulfoxide (DMSO) were supplied by Sigma Aldrich, Canada. Fluo-4 AM calcium indicator (#F14217), 4′,6-aiamidino-2-phenylindole dihydrochloride (DAPI, #D1306), Probenecid (#P36400), cardiac Troponin T (cTnT) monoclonal antibody (#MA5-12960), and secondary antibodies (including goat anti-Mouse Alexa Fluor 488 (#A11001), goat anti-rabbit Alexa Fluor 647 (#A21244), and goat anti-mouse Alexa Fluor 647 (#A21235)) were purchased from Invitrogen, USA. Tyrode’s buffer was obtained from Alfa Aesar, USA. Phalloidin-iFluor 488 reagent (#ab176753) and recombinant Anti-Vimentin antibody (#ab92547) were obtained from Abcam, UK.

### 2.2. Chip Design

The microphysiological devices, commercially known as OMEGA^MP^, were supplied by eNUVIO Inc. (Montréal, QC, Canada). These chips were fabricated from PDMS and featured two cell-seeding chambers. Each chamber contained an array of micropillar pairs designed to support tissue attachment and anchorage. With a diameter of 21.25 mm, each device is specifically designed to fit tightly into the wells of standard 12-well culture plates, facilitating seamless integration into existing laboratory workflows. As schematically illustrated in **Fig. S1**, the seeding chamber has an elliptical shape (8 mm × 4 mm × 2 mm), providing a surface area of 0.25 cm² and a maximum chamber volume of 40 μl. The pillars within the chamber are 3 mm in height and feature a tilted design, effectively preventing the tissue from slipping off the pillars and ensuring stable tissue anchorage. The pillar diameter was designed to taper as the distance from the PDMS base increased, with an average diameter of 1 mm. The PDMS base beneath the seeding chambers has a thickness of 0.2 mm, providing a thin and transparent layer that facilitates high-quality microscopic readouts directly on the chip.

### 2.3. eMSG Fabrication

eMSGs were fabricated using a two-phase emulsion system comprising an aqueous phase and an oil phase. According to the previous work, this method allowed for the formation of uniform, cell-sized hydrogel droplets with well-defined mechanical properties suitable for embedding in 3D tissues [24]. The oil phase consisted of a 6% (w/v) solution of PGPR 4150 in kerosene. The aqueous phase comprised a very soft polyacrylamide pre-polymer solution (E ∼ 0.15 kPa) containing 0.2 μm carboxylate-modified polystyrene nanoparticles for fluorescent edge labeling, which was visible at Texas Red wavelengths (excitation peak at 586 nm and emission peak at 603 nm). All fabrication steps were conducted inside a fume hood, and glass tubes were used as containers for all solutions to ensure a controlled and contamination-free environment. Prior to mixing, all solutions were purged with nitrogen gas for 20 min to eliminate any oxygen in the liquid phase. To initiate crosslinking, a solution containing 1 wt% ammonium persulfate (APS) was added to the pre-polymer solution at a 1:10 ratio. This mixture was then immediately transferred to the oil phase. The combined phases were vortexed for 7 seconds at 3000 rpm and subsequently stirred under oxygen-free conditions for 20 minutes to allow the gelation process to complete and form hydrogel beads. After stirring, the mixture was left undisturbed for 20–30 minutes, enabling the hydrogel beads to settle at the bottom of the tube by gravity. The PGPR was removed by washing the hydrogel beads with fresh kerosene multiple times until the liquid became completely clear, ensuring the removal of residual surfactant and oil phase components. To remove kerosene, the hydrogel beads were washed several times with PBS. After each wash, the beads were collected by centrifugation at 21,000 g for 3 minutes. Once cleaned, the surface of the fabricated beads was functionalized with bovine collagen type I to enhance their biocompatibility and facilitate cellular interactions. Briefly, after overnight UV sterilization, the collected beads were resuspended in a solution of 0.1 mg/mL Sulfo-SANPAH in PBS and exposed to UV light for 4 minutes to facilitate the formation of crosslinking radicals. After removing Sulfo-SANPAH via centrifugation at 21,000 g for 3 minutes, the beads were resuspended in a 0.1 mg/mL solution of bovine collagen type I and incubated overnight at 4°C to ensure thorough functionalization. The following day, the collagen solution was removed by washing the beads three times with PBS. The sterile, collagen-functionalized beads were then stored at 4°C for future use.

### 2.4. Cardiac Cell Isolation and Culture

Primary ventricular CMs and cardiac fibroblasts (CFs) were isolated from neonatal Sprague-Dawley rats (Charles River Laboratories, Canada) using a Neonatal Heart Dissociation Kit (Miltenyi Biotec, Germany), following the protocol described in our previous reports [31, 32]. The procedure was conducted in accordance with the protocol approved by the Animal Ethics Committee of the Centre Hospitalier Universitaire Sainte-Justine Research Centre (CRCHUSJ, protocol number: 2023-4833) and adhered to the guidelines of the Canadian Council on Animal Care (CCAC) and the US National Institutes of Health Guide for the Care and Use of Laboratory Animals. Briefly, 2-day-old Sprague-Dawley rat neonates were euthanized by decapitation, and their hearts were immediately collected in cold PBS. The ventricles were dissected into small pieces and washed twice with cold PBS. Heart dissociation was performed using a combination of mechanical and enzymatic digestion, as per the manufacturer’s protocol, with the gentleMACS Dissociator (Miltenyi Biotec, Germany). Following dissociation, the tissue was treated with a red blood cell (RBC) lysis solution (Miltenyi Biotec, Germany) following the manufacturer’s instructions. The dissociated cells were pre-plated in DMEM/F12 culture medium, supplemented with 10% FBS and 1% Pen Strep. After 1 hour, CMs that remained suspended in the medium were separated from CFs, which adhered to the flask. The suspended CMs were collected as cell pellets by centrifugation at 600 × g for 5 minutes. Adherent CFs were maintained in the DMEM/F12 complete medium mentioned above and used at passage 2 (P2) upon reaching 80% confluency. Primary CMs were immediately used for cell encapsulation by resuspending the cell pellets in the prepared hydrogel. Before encapsulation, the isolated CMs were counted, and their viability was assessed using a trypan blue assay with dual-chamber cell counting slides (Bio-Rad Laboratories, Canada) and the Countess II FL automated cell counting machine (Thermo Fisher Scientific, USA).

### 2.5. Engineered Heart Tissue (EHT) Formation

EHTs were created by encapsulating isolated CMs in a Fibrin/Geltrex hydrogel and seeding the mixture into each chamber of the fabricated chip. To ensure sterility during cell culture, all prepared solutions were sterilized using 0.2 μm polyethersulfone (PES) syringe filters. Before cell seeding, the sterile chips were carefully transferred into individual wells of 12-well plates using tweezers under sterile conditions. The device was coated by adding 2 mL of a 5% Pluronic F-127 solution (prepared in PBS) to each well and incubating it overnight at 4°C. This antifouling coating was applied to prevent cell adhesion to the base and walls of the PDMS chambers. All subsequent procedures for tissue formation were conducted on ice, using pre-cooled solutions and pipette tips to maintain optimal conditions.

The isolated CM pellet was resuspended in a fibrinogen solution. Geltrex matrix and the eMSG solutions were then added to the cell-hydrogel mixture to create the final suspension. At this stage, the Pluronic F-127 coating solution was aspirated from the microplate chips, and each well was washed once with cold PBS. After washing, the PBS was completely removed from the chambers to prepare them for cell-hydrogel seeding. Next, thrombin solution was added to the cell-hydrogel mixture, mixed thoroughly and quickly, and immediately dispensed over the entire seeding chamber to ensure uniform distribution without introducing air bubbles. A seeding volume of 30 μL was used for each individual microtissue, with each containing approximately 450,000 cells (equivalent to a cell density of 15 million cells/ml). The plates were then incubated for 15 minutes to allow complete fibrin polymerization and gelation. After gelation, the microtissues were cultured in Claycomb medium supplemented with 10% FBS, 1% Pen Strep, 1% GlutaMAX, and 10 μg/mL aprotinin. To fully immerse the device (containing two microtissues per chip), 2 mL of culture medium was added per well. The culture medium was replaced the next day and subsequently refreshed every other day to maintain optimal conditions for tissue growth and development. Three different fibrinogen concentrations were used in this study to create microtissues with final fibrinogen concentrations of 2.5, 5, and 10 mg/mL, forming F2.5, F5, and F10 cardiac microtissues, respectively. Across all experimental groups, the final concentration of Geltrex was fixed at 20% v/v, and the thrombin concentration was standardized at 0.2 Units/mg of fibrinogen. Tissue compaction was monitored over 14 days of culture using a Leica DMi8 wide-field microscope (Leica Biosystems, Germany). As a control group for tissue compaction analysis, CFs were seeded at the same cell density in the F5 microtissue to evaluate the impact of cell type on tissue compaction.

### 2.6. Immunofluorescence (IF) Staining

The cardiac microtissues with varying fibrin concentrations were stained for cardiac-specific markers, as described in our previous report [31]. EHTs were fixed in 4% PFA solution for 20 minutes at room temperature (RT). The microtissues were then permeabilized with 0.3% Triton-X 100 in DPBS for 30 minutes and subsequently incubated in a blocking buffer containing 5% BSA and 0.3% Triton-X 100 in DPBS at RT for 1 hour. An anti-vimentin antibody was used for CFs, while a cTnT antibody was used for CMs. The EHTs were incubated with the primary antibody solution (1:200) diluted in the blocking buffer at 4°C overnight. The microtissues were then incubated in a blocking buffer containing secondary antibodies (1:400) and phalloidin (1:1000) at RT for 1 hour. Nuclei were counterstained using DAPI solution (1:1000) for 20 minutes. Finally, the EHTs were washed three times with DPBS, and confocal imaging was conducted using the SP8-DLS Leica microscope (Leica Biosystems, Germany).

### 2.7. Pillar Mechanical Properties

The force-displacement relationship of PDMS pillars was evaluated by measuring the force required to deflect the pillars using a MicroSquisher device (CellScale, Canada), as described previously [33]. Briefly, a circular tungsten microbeam with a diameter of 558.8 μm was used to apply the displacement. Before testing, the platform was securely fixed on the testing stage. The probe tip was positioned at various distances from the pillar base—500, 800, 1000, and 1500 µm. The probe applied force perpendicularly to the pillar’s orientation, displacing it by 200 µm over 40 seconds. The force was then held constant for 2 seconds before being released, allowing the pillar to return to its original position within 40 seconds. Three pillars were tested at each specified distance. The resulting experimental data were fitted to a linear equation to generate a force-displacement curve, averaged across the three pillars. The slope of this calibration curve was then used to calculate the contractile force at each specific tissue position. The relationship between this coefficient and the distance from the pillar base was determined by fitting the experimental data from the four tested positions to an exponential equation. Additionally, videos were created using Microsoft Clipchamp, combining images of the pillar captured during the process.

### 2.8. Calcium Imaging

Calcium transients were assessed using the Fluo-4 AM calcium indicator assay, as described previously [34]. After 7 days of culture, the cardiac microtissues were incubated in Tyrode’s buffer supplemented with 5 μM Fluo-4 AM, 1 mM Probenecid, and 0.025% Pluronic F127 at 37°C for 45 minutes. The samples were then transferred to fresh Tyrode’s buffer at 37°C for 30 minutes to equilibrate. Calcium transient videos were recorded using a Leica DMi8 fluorescent microscope equipped with a GFP filter cube, operating at a fast-readout rate of 50 frames per second (fps). The fluorescent videos were analyzed using FIJI software (NIH, USA), where the fluorescence intensity (F) of a defined region of interest (ROI) was calculated and plotted over time, normalized to the background fluorescence intensity (F0) [35].

### 2.9. Tissue-scale Contractility

Bright-field videos of the contracting EHTs were recorded on-chip using a Leica DMI8 inverted microscope equipped with a 5x objective. The videos were analyzed using an iterative particle image velocimetry (PIV) method in MATLAB R2023b to track the movement of the pillars over time, as previously described [36]. The contractile force was calculated using the previously mentioned pillar force-displacement linear equation described in Section 2.7. The calculated force was further normalized by dividing it by the cross-sectional area of each tissue.

### 2.10. Pharmacological Studies

Two drug candidates with known effects were used for drug testing: Norepinephrine (NE) and Blebbistatin (Bleb). Both compounds were dissolved in DMSO according to the manufacturer’s recommended protocol and further diluted in culture media to a final concentration of 100 μM. First, NE solution was added to the cardiac microtissues, followed by 30 minutes of incubation at 37°C. The beating and contraction of the cardiac microtissues were recorded before and after the addition of NE using the previously described method. Subsequently, the samples were washed three times with fresh media, with a 5-minute incubation at 37°C after each media change. Afterward, Bleb solution was added to the samples, and the contractile activity was recorded again after 30 minutes of incubation at 37°C [37].

### 2.11. Cell-scale Stress Measurements

Cell-scale internal stresses were calculated based on eMSG deformations, as previously reported [24]. The eMSG images were recorded using a DMi8 fluorescent microscope equipped with a TXR filter cube and a 20x objective. The sensor shape was measured with FIJI software at different time points (Day 0, 3, 7, and 14). Points along the edges of each bead were selected, and the shapes were fitted as circles under stress-free conditions (Day 0) or as ellipses under stressed conditions (Days 3, 7, and 14). The lateral and longitudinal orientations were defined based on the tissue formation shape. The longitudinal orientation corresponded to the direction of tissue alignment between two pillars (x-axis), while the lateral orientation was defined as the direction of tissue width compaction (y-axis), as illustrated in **Fig. 7B**. Longitudinal and lateral strains were calculated based on the sensor deformation and size changes in the x and y dimensions. A custom-built MATLAB code was developed to convert these local strains into stress values. To account for measurement error, the sensor size was measured three times, providing the mean and standard deviation of measurements in both orientations. A Monte Carlo-based analysis was conducted, generating 10,000 randomly produced strain values following a Gaussian probability distribution. This distribution was then used to compute the 95% confidence intervals for each data point, indicating the upper and lower limits of the corresponding stress values.

### 2.12. Finite Element Analysis for Stress Measurements in 3D

COMSOL Multiphysics v6.2 was used to create a 3D model of the eMSGs to evaluate the impact of additional deformations of 10–20% strain in the z-axis (along the height of the tissue). The hyperelastic material properties were applied as previously described [25] with Neo-Hookean Lamé parameters: shear modulus *G* = 60 Pa, Poisson ratio *ν* = 0.3, *Lamé 1* = 60 Pa, and *Lamé 2* = 90 Pa). The eMSG geometry was initially modeled as spherical, with strain boundary conditions applied to the solid domain in the x, y, and z axes based on experimentally relevant values. Principal components of the stress tensor were evaluated along each axis to determine stresses in 3D. A triple parametric sweep was conducted, varying strains in the x, y, and z axes and probing the principal stresses along each respective axis to generate stress variation curves under increasing compressive strains in the z-axis. The results showed that stresses in the x and y axes were underestimated by approximately 15–75 Pa when larger compressive strains were applied in the z-axis.

### 2.13. Sensor Deformation during Contraction

The sensor deformation during beating was measured through image processing of the recorded fluorescent videos using MATLAB R2021a software [38]. Briefly, the contours of the beating shape across various time frames were tracked by processing each frame of the data series. The analysis began by converting each frame from RGB to grayscale to facilitate a more detailed examination. For each frame, a region of interest (ROI) was carefully defined, with the corresponding grayscale image displayed for clarity. A global threshold value of 28 was set to binarize the images. This process categorized pixels with intensities at or above 28 as binary ‘1’s, while all others were assigned a value of ‘0’. Although the initial contour detection provided a general outline of the shape, it was sensitive to image discontinuities and the presence of holes. To address these issues, morphological operations were applied to the binarized images. These operations included filling voids, removing small artifacts, and smoothing edges, which ensured the extraction and presentation of the most prominent contour. This contour was then transformed into a mask, and its pixel area was calculated and converted into a physical area using a predefined scale factor. The results were presented as area data across different frames, showing the temporal evolution of the beating shape’s contour. Changes in the shape were subsequently translated into cell-generated stress variations using the MATLAB code described in Section 2.11, providing a comprehensive assessment of the beating and contracting behavior of the tissue.

### 2.14. Statistical Analysis

Statistical analysis was performed using GraphPad Prism software version 9.0 (GraphPad Inc., San Diego, CA, USA). Appropriate statistical tests were selected based on the number of experimental groups. One-way analysis of variance (ANOVA) followed by Tukey’s post hoc test was applied when there were three or more study groups, while unpaired t-tests were used for comparisons between two groups. Unless otherwise specified, all experiments were conducted with three independent replicates. Data are presented as the mean ± S.D. A p-value of less than 0.05 was considered statistically significant.

## 3. Results and Discussion

### 3.1. Study Overview

We report the application of a PDMS-based HOC model to fabricate micro-scale beating cardiac tissues by encapsulating primary cardiac cells in a fibrin/Geltrex matrix. The cell-laden hydrogels compacted between two elastic pillars which serve as anchorage for tissue formation and can be used to calculate contractile forces based on their mechanical properties. In addition to global contractility measurements at the tissue scale via pillar deflection, the integration of ultrasoft mechanosensors allows for continuous, local, cell-scale measurements. The transparency of the PDMS chip enables real-time monitoring of tissue contractile properties based on pillar deflection (via bright-field imaging), cell-generated stresses based on eMSG deformation, and intracellular calcium transients (via multichannel fluorescent imaging). Real-time imaging of contractile activity is particularly important for drug screening applications to study the inotropic and chronotropic effects of pharmaceutical compounds. In addition to these live-tissue experiments, our HOC model also demonstrated suitability for other tissue-based investigations (e.g., IF staining) as well (**Fig. 1**).

**Fig. 1.**
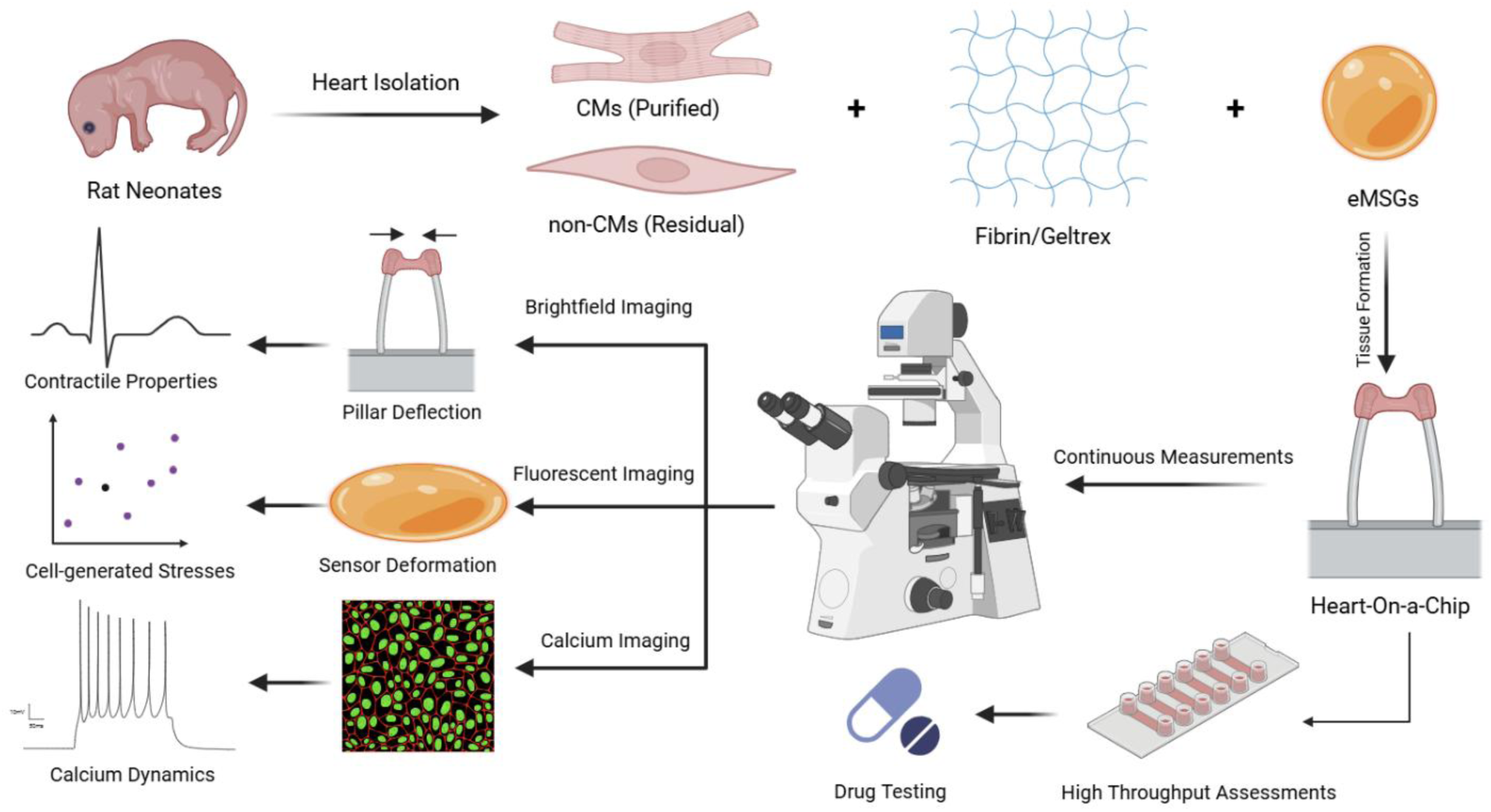
A schematic representation of the study describes tissue formation by encapsulating neonatal rat cardiac cells within a fibrin/Geltrex hydrogel mixture integrated with eMSGs, continuous measurement of global contractile properties based on pillar deflection, local cell-generated stresses based on sensor deformation, calcium handling properties, and high-throughput assessments in 12-well plates for drug screening applications (produced by BioRender).

### 3.2. Tissue Formation and Compaction

Cardiac microtissues were developed by encapsulating purified primary CMs into a fibrin/Geltrex hydrogel and seeding them into the chamber of the device. Fibrin is a naturally occurring biopolymer that plays a key role in the wound-healing process by stabilizing the structure of blood clots. Fibrin can be immediately formed by mixing fibrinogen and thrombin and degraded in just a few hours following polymerization in vivo through a process called fibrinolysis [39, 40]. Fibrin also demonstrates rapid in vitro degradation due to the presence of plasmin and non-plasmin proteases in the cell culture medium, which are secreted by the cultivated cells. Therefore, for in vitro studies using fibrin hydrogel, a fibrinolysis inhibitor (e.g., aminocaproic acid, tranexamic acid, aprotinin) is used to enhance the stability of fibrin [41].

In this study, the cardiac microtissues were cultured in a specialized medium called Claycomb medium, which has been previously shown to maintain the contractility of primary CMs [42]. Aprotinin was added to the culture medium as a fibrinolysis inhibitor, which is necessary for tissue formation and stability, as well as matrix deposition and remodeling. The selection of aprotinin was based on several previous reports of its use as a fibrinolytic inhibitor in cardiac tissue engineering applications [41–43]. Additionally, compared to other stabilizing agents, aprotinin inhibits a broader range of serine proteases [44]. Geltrex is another component of our hydrogel; it is a basement membrane extract, mainly composed of laminin, collagen IV, entactin, and heparan sulfate [45]. This component was added to the fibrin hydrogel to recapitulate the ECM composition for the encapsulated cardiac cells by providing several important proteins and proteoglycans.

The platform was incorporated into 12-well plates and was shown to support reproducible tissue formation and compaction (**Fig. 2A**). The minimal volume of the cell-hydrogel mixture required for the formation of a uniform tissue was used for cell seeding, which was experimentally determined to be 30 μl. This strategy minimized the tissue size and the number of cells needed per tissue. The cell-laden hydrogel formed immediately after the addition of thrombin to the system and was seeded into each chamber. The cells significantly remodeled their matrix during the first three days of culture and generated a dense tissue suspended between two pillars. Tissue compaction gradually continued over two weeks (**Fig. 2B**).

**Fig. 2.**
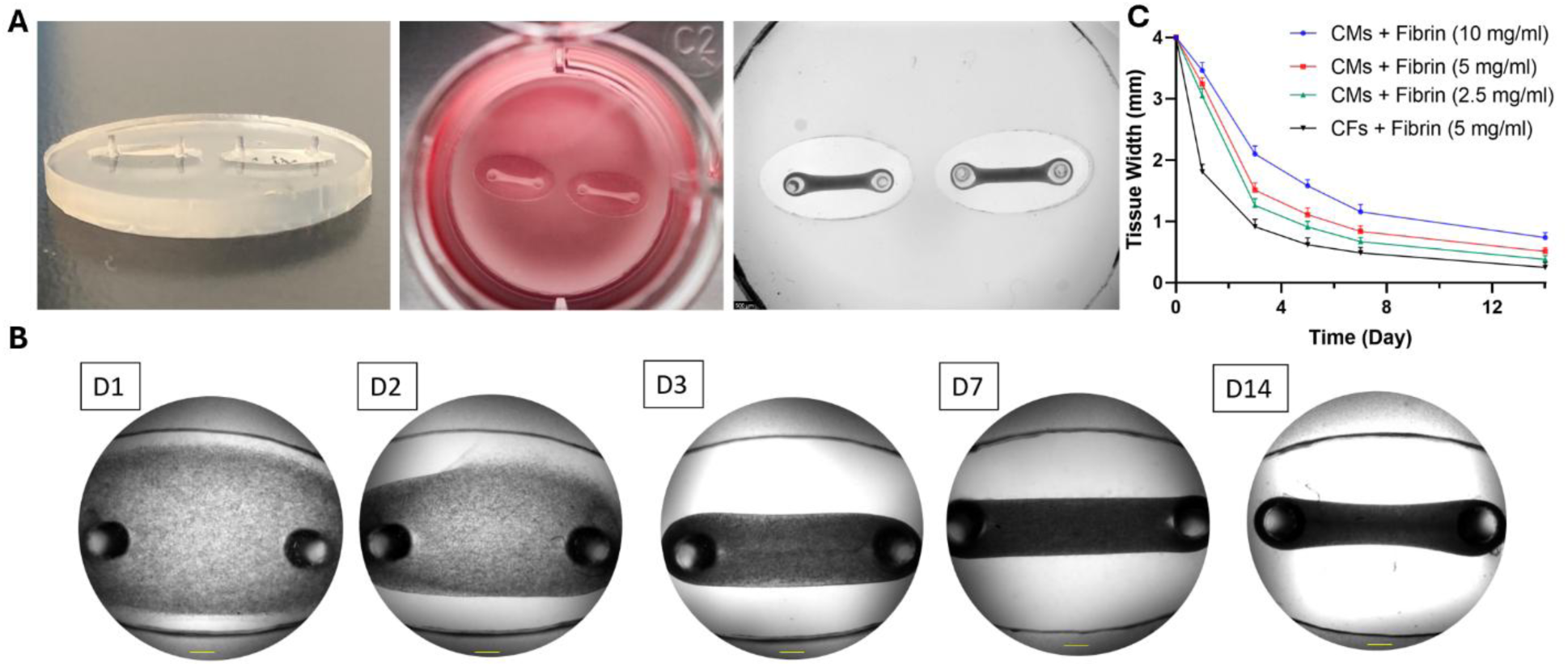
Tissue Formation and Compaction. (A) Demonstration of the chip, including a side view, a top view, and reproducible tissue formation. (B) Gradual tissue compaction over two weeks of culture. Scale bar = 500 μm. (C) Effects of fibrin concentration and cell type on tissue compaction.

The elliptical shape of the chambers supported the formation of EHTs as tissue strips without the formation of a necking area in the middle of the tissue, which typically leads to tissue failure and breakage due to higher residual stress in the necking region, as previously reported for rectangular molds [46, 47]. Therefore, long-term culture of cardiac microtissues could be achieved in our system without failure. Although higher tissue compaction is generally known to be a crucial factor for the generation of functional cardiac tissues, extremely high tissue compaction leads to early tissue degradation and failure [20, 48]. This unfavorable remodeling could result from very low polymer concentration, very high cell density, or a high ratio of non-CMs in the cardiac cell blend [21]. Therefore, several factors, such as matrix concentration, stiffness, cell density, and the ratio of CMs to non-CMs, should be carefully optimized to generate a beating cardiac microtissue [49].

In this study, three different fibrin concentrations (2.5, 5, and 10 mg/ml) were used to assess the effect of matrix concentration and stiffness on tissue compaction. As shown in **Fig. 2C**, increasing the fibrin concentration resulted in reduced tissue compaction. The tissue width was significantly lower than the initial seeding value (4 mm in the middle of the tissue) in all experimental groups. After 7 days of culture, the width of the tissue was reduced to 29%, 21%, and 16.7% of the initial value for F10, F5, and F2.5 EHTs, respectively. Therefore, a shift from milli-scale to micro-scale cardiac tissues was observed by adjusting the polymer concentration, and the width of all tissues became less than 1 mm after 14 days of culture. This micro-scale tissue is favorable for in vitro studies because it requires a lower number of cells, materials, and reagents while maintaining adequate mass for structural, functional, and molecular experiments with convenient handling [50].

The reduced compaction in more concentrated hydrogels could be due to the fact that higher polymer concentration leads to a denser and stiffer network of fibrin fibers with smaller pore sizes, which impedes cell migration, interaction, and their ability to exert traction forces [51]. Therefore, the more concentrated matrix resists deformation and remodeling to a greater extent, which leads to lower and more delayed tissue compaction [52]. As a control, CFs were encapsulated in a fibrin (5 mg/ml) hydrogel and seeded at the same density and volume. Tissue compaction became significantly faster and more pronounced in this CF control compared to all the CM tissues, and the tissue strip was formed within one day. This could be correlated with the known ability of fibroblasts to actively produce and remodel the ECM. However, CMs are the contractile machinery of the heart, and their capacity for migration and matrix compaction is much more limited [53–55]. It is worth mentioning that the cardiac microtissues were generated using purified neonatal rat CMs, and a portion of non-CMs was still included during cell encapsulation and tissue formation. Thus, the compaction of the CM microtissues could be attributed to the low ratio of non-CMs.

### 3.3. Immunohistochemistry of Cardiac Microtissues

An IF assay was performed on F2.5, F5, and F10 cardiac microtissues after 7 days of culture by co-staining with DAPI, phalloidin, and cTnT to visualize the structure and morphology of the encapsulated CMs. The confocal images are shown in **Fig. 3A-C** for F2.5, F5, and F10 samples, respectively, which represent the overall structure of the tissues as well as higher magnifications under the 10x objective (**Fig. S2-3**). Microscopic images of these EHTs at other magnifications (5x and 20x) are provided in **Fig. S4-5**. DAPI and phalloidin label the cell nuclei and cytoskeletal F-actin protein, respectively. Additionally, cTnT, a structural protein, is a key component of the troponin complex and plays an important role in cardiac muscle contraction. It is also a cardiac-specific biomarker and can be selectively used for the staining and imaging of CMs [33]. As shown in **Fig. 3**, whole-mount staining of all three cardiac microtissues revealed the presence of aligned CMs with an elongated structure, expressing cardiac-specific biomarkers. This cell alignment is critical for the spontaneous and synchronous beating of EHTs. Higher-resolution images indicated the relevant phenotype of CMs with a rod-shaped morphology interconnected to each other, displaying elongated and striated fibers. The constructs were highly compacted, containing a large number of cells; however, lower compaction was easily observed in higher fibrin concentrations. Additionally, relatively more rounded cardiac cells were observed in higher fibrin concentrations (F10 microtissue), which could be due to the higher matrix stiffness. In contrast, more compacted and interconnected CM networks with highly aligned and elongated cells were observed in lower fibrin concentrations (F5 and F2.5 microtissues). Higher cell spreading in lower fibrin concentrations has been previously reported, which is consistent with our findings [48, 56].

**Fig. 3.**
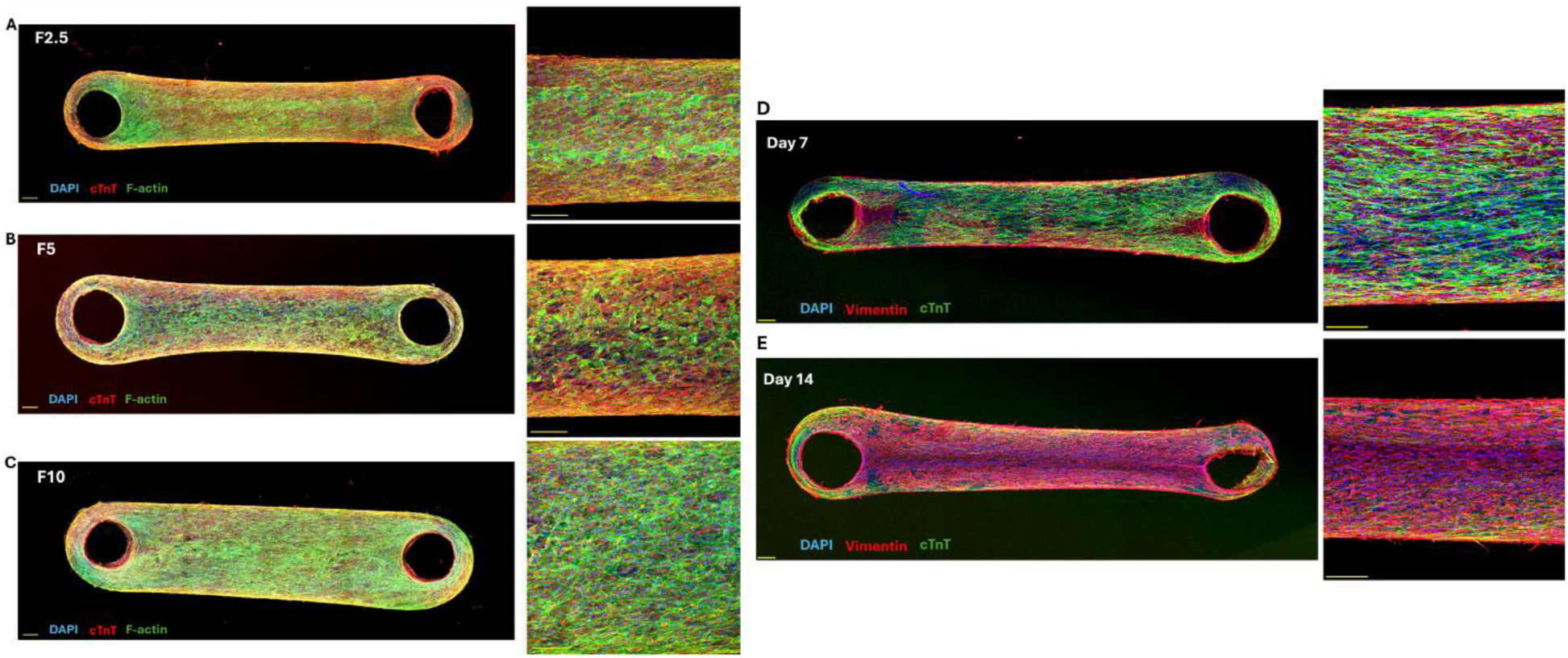
Confocal Imaging. Whole-mount IF staining as well as higher-resolution confocal images of (A) F2.5, (B) F5, and (C) F10 cardiac microtissues after 7 days of culture. Merged images of DAPI (blue), cTnT (red), and F-actin (green). Representative confocal images of cardiac microtissues after (D) 1 week and (E) 2 weeks of culture. Merged images of DAPI (blue), cTnT (green), and vimentin (red). Scale bar = 200 µm.

Since the CM purification via preplating method do not result in 100% pure CMs, a portion of non-CMs may remain in the final cell pellet. To identify non-CMs in our EHTs, vimentin was stained, which is an intermediate filamentous protein that maintains the structural integrity and stability of the cytoplasm. IF staining of this biomarker has been previously reported for imaging non-CMs in cardiovascular studies [58]. In addition, IF staining of cell nuclei and CMs was carried out with DAPI and cTnT, respectively, as explained previously. On day 7 of culture, the striated and elongated fibers of CMs were observed with a high level of alignment and interconnection. Non-CMs also showed elongated spindle-like morphology, forming a network mainly on the peripheries of the tissue (**Fig. 3D and S6**). The spindle-like morphology is consistent with the phenotype of a wide range of non-CMs (e.g., CFs and endothelial cells). Furthermore, the dominance of CMs over non-CMs was evident based on the microscopic images, indicating that most of the tissue area was covered with CMs. On the other hand, non-CMs dominated the tissue by covering a major part of it on day 14 as the constructs continued to become more compacted (**Fig. 3E and S7**). Although the CMs partially retained their elongation and alignment on the edge of the tissue, their interconnection was disrupted by a large network of non-CMs. This prevalence of non-CMs after two weeks of culture could be explained by their high ability to grow and proliferate, unlike CMs, which have limited proliferation capacity [59, 60]. Therefore, the number of non-CMs significantly increased over time compared to CMs, which adversely affected the alignment, elongation, interconnection, and function of the CM network in cardiac microtissues.

### 3.4. Mechanical Properties of Pillars

The flexible pillars provide anchor points for tissue attachment and maintain passive tension during auxotonic tissue contraction. The deflection of the pillars can be directly used to calculate contractile forces based on the bending beam equation [61]. The coefficient of the pillar force-displacement relationship depends on several factors, such as pillar mechanical properties (e.g., Young’s modulus), pillar diameter, and pillar length [10]. Various factors also regulate the elastic modulus of the PDMS pillars, including the curing temperature, curing duration, and the ratio of the curing agent to the elastomer [50]. Here, we determined this calibration curve based on Microsquisher analysis, and the linear equation between the applied force and the detected deflection was directly obtained. Three different pillars were investigated as independent replicates, and the test was performed on four different pillar heights (distance from the PDMS base) by placing the Microsquisher probe on each pillar height (**Movie S1-4**).

The force-displacement experimental data for different pillar heights (500, 800, 1000, and 1500 μm) are depicted in **Fig. 4A**. Increasing the pillar height (i.e., being more distant from the base) reduced the slope of the force-displacement curve, which was consistently observed for each pillar. This means that applying the same amount of force to a higher pillar height results in greater pillar deflection than with a lower pillar height. This trend has been computationally modeled in a previous study [33]. Additionally, the alteration of this slope by changing the pillar height was not linear, likely due to the continuous reduction in pillar thickness as the height increased.

**Fig. 4.**
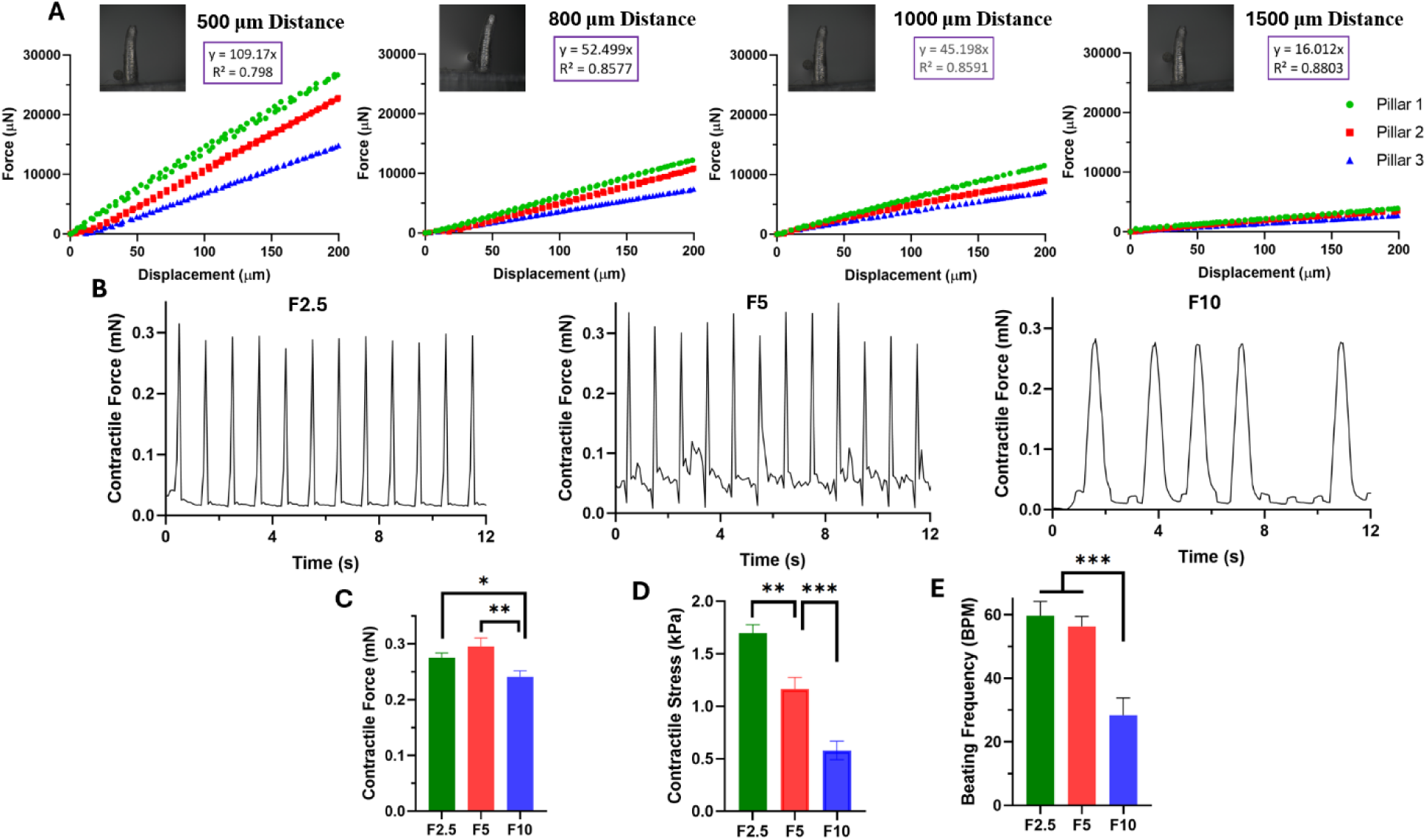
Contractile Properties. (A) Force-displacement relationship of pillars at different heights. (B) Beating patterns of EHTs with different fibrin concentrations, which were quantified for (C) contractile force, (D) contractile stress, and (E) beating frequency (*, **, *** show statistical significance at p<0.05, p<0.01, and p<0.001, respectively).

At each specific height, there were some variations between the independent replicates (i.e., three tested pillars), and these variations consistently decreased at higher pillar heights. However, the average linear equation was used for each height since the correlation coefficient (R²) was acceptable for all of them (0.8 or higher). The slope values of these linear regressions were 109.170, 52.499, 45.198, and 16.012 N/m for the lowest to highest pillar height, respectively. These values can be further used to calculate tissue contractile forces based on the tissue position. Additionally, no hysteresis was observed for these pillars. Therefore, the same coefficient could be used to determine both tissue contraction and relaxation properties. Moreover, the force-displacement coefficients were fitted versus pillar height based on these experimental values, and an exponential formula was derived to estimate this coefficient for other pillar heights (**Fig. S8**).

As expected, there were some batch-to-batch differences in the mechanical properties of the pillars. The ideal condition is to perform a series of experiments on a single platform that has been recently manufactured and fully characterized. However, linear regression (with an appropriate correlation coefficient) has been reported in previous literature as an acceptable estimation for the force-displacement relationship of pillars [62, 63].

### 3.5. Functional Assessments

The functional properties of cardiac microtissues were further evaluated based on contractile activity, calcium transients, and pharmacological intervention. Following tissue formation, the constructs were monitored daily for tissue contraction. After the constructs became compacted on day 3 and the tissue strip formed, the EHTs began beating spontaneously. However, the tissue contraction was not forceful enough to move the pillars on the third day of culture, and the pillar deflection was negligible. The tissue contraction became more vigorous over time, reaching a peak on day 7. Afterward, the EHTs stopped contracting. This beating trend is consistent with previous reports on neonatal rat CMs [64, 65].

#### 3.5.1. Contractile Properties

The contractility of F2.5, F5, and F10 cardiac microtissues was recorded using an inverted bright-field microscope (**Movie S5-7**). The contractile force is proportional to the displacement of the pillars and can be calculated based on the force-displacement relationship of the pillars at a specific tissue position. The tissues mostly formed in the range of 800 to 1000 μm above the chamber base. This means that the EHTs were placed in the lower one-third of the total pillar height, which helps maintain the tissues on the chip during beating without removal or tissue failure.

**Fig. 4B** demonstrates the beating pattern of different EHTs over time, which was quantified for contractile force (mN) in **Fig. 4C**. These values were further normalized to the cross-sectional area of the EHTs in **Fig. 4D** to obtain the contractile stress (mN/mm²). This normalization allows for universal comparison to other HOC platforms and is also required by regulatory agencies and different consortia [33]. The beating frequency is also indicated in **Fig. 4E** as beats per minute (BPM).

The beating patterns showed higher beating frequency, beating regularity, and contractile forces at lower fibrin concentrations. The amount of generated contractile force ranged from 0.2 to 0.3 mN for the different tissues. The differences in contractile force were not significant between the F2.5 and F5 samples, while both were significantly higher than the F10 group. The trend in contractile force was consistent with the previous report by the Tranquillo group [66]. The highest contractile stress was observed in the most compacted tissue with the smallest cross-sectional area. The contractile stress of F2.5 microtissues was approximately 1.45- and 2.92-fold higher than that of the F5 and F10 groups, respectively, with values ranging from 0.5 to 1.8 mN/mm². Our contractile values are within the range of those previously developed for EHTs by the Eschenhagen group. However, differences in cell density, seeding volume, tissue species, matrix composition, and external stimulation should be considered [67]. The beating frequency followed the same trend as contractile force, with no significant difference between the F2.5 and F5 groups, while both were approximately 2-fold higher than the F10 sample. The contractile frequency values ranged from 25 to 65 BPM. Higher fibrin concentrations also resulted in greater beating irregularities in terms of frequency and amplitude over time.

Overall, we observed diminished contractile activity at higher fibrin concentrations, especially when increased to 10 mg/ml. This weakened contractility could be attributed to several factors. Higher polymer concentration generally leads to higher matrix stiffness, which can provide structural stability for the encapsulated cells. However, excessive stiffness may impair cell-cell and cell-ECM interactions, reduce cell spreading and alignment, disrupt intercellular gap junctions and electromechanical coupling, decrease cell-mediated compaction, and limit oxygen and nutrient transport within the denser network. It could also hinder the ability of CMs to complete their contraction-relaxation cycle. In this study, the excessive fibrin concentration was 10 mg/ml, which impaired the function of the encapsulated CMs. However, increasing the fibrin concentration from 2.5 to 5 mg/ml did not adversely affect the activity of the EHTs. Therefore, it can be concluded that using fibrin hydrogel with a concentration as high as 5 mg/ml could support the functionality of cardiac cells while still providing their structural stability, which is important for long-term cell culture.

#### 3.5.2. Calcium Handling

Calcium handling of the EHTs was evaluated using non-invasive fluorescent-based imaging. Calcium mobilization between adjacent CMs is based on the difference in intracellular calcium ion concentration, which plays an important role in the process of calcium-induced calcium release and leads to the contraction-relaxation coupling of CMs [31].

Therefore, evaluating the calcium transient profile of fabricated tissues is crucial for identifying the optimal conditions. Here, we investigated the effect of matrix composition on the calcium handling of EHTs by assessing two different experimental groups (i.e., F5 and F10) with low and high fibrin concentrations, respectively. The calcium dynamics were recorded on day 7 using fast, high-resolution microscopy and are shown in **Movies S8-9** for F5 and F10 samples, respectively. On day 14, the tissues did not show the contractile profile, and calcium imaging indicated a lack of intracellular calcium mobilization (**Movie S10**). The beating cardiac syncytium, with visible calcium wave propagation, was observed in both EHTs on day 7. However, a more synchronous uniaxial longitudinal conduction was observed at lower fibrin concentration. At higher fibrin concentration, high magnification of calcium traces revealed an evident irregularity in the calcium transient of different regions of the EHTs. The visual videos were further displayed in patterns based on the change in fluorescence intensity over time and are shown in **Fig. 5A-B** for F5 and F10 EHTs, respectively. Higher beating regularity, synchrony, and frequency were observed at lower fibrin concentration, which is consistent with our contractility results. The calcium handling parameters, including amplitude, time to peak, and relaxation time (from peak to baseline (90%)), were further quantified based on this graph (**Fig. 5C-E**). Lower fibrin concentration resulted in significantly higher calcium transient amplitude, faster calcium release, and more rapid calcium re-uptake. Prolonged calcium handling capacity with wider waves and more irregular peaks was observed at higher fibrin concentration, indicating the relative immaturity of sarcoplasmic reticular function in these EHTs. This impaired calcium signaling could be attributed to the lower interconnection of CMs in the denser polymeric networks, which aligns with our contractility results and discussion.

**Fig. 5.**
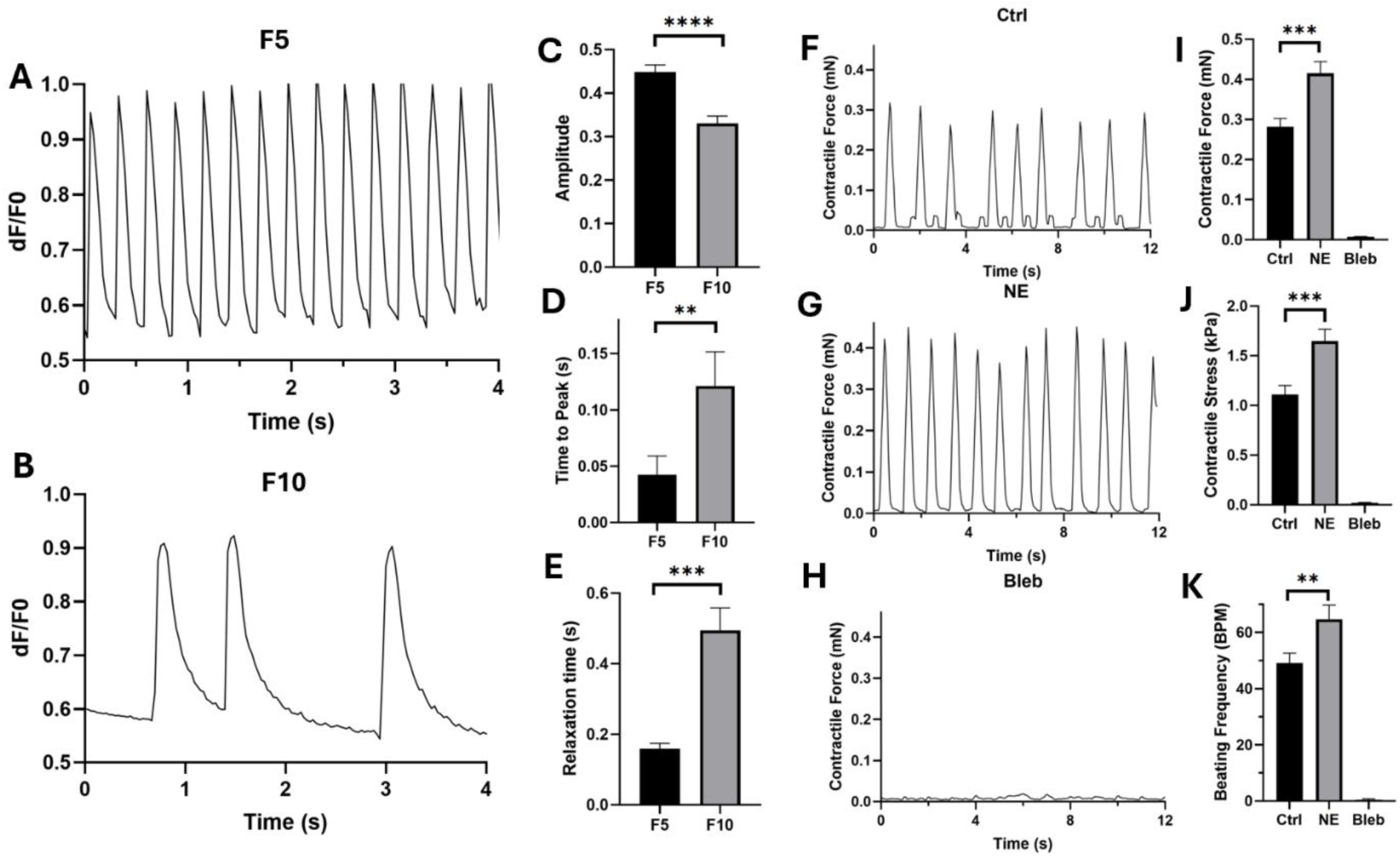
Calcium Handling and Drug Testing. Calcium transient graphs representative of the change in intracellular calcium concentration for F5 (A) and F10 (B) microtissues, which were quantified for (C) amplitude, (D) time to peak, and (E) relaxation time. Beating patterns of the control tissue (F) and drug-treated tissue with norepinephrine (NE) (G) and blebbistatin (Bleb) (H), which were quantified for (I) contractile force, (J) contractile stress, and (K) beating frequency (**, ***, **** indicate statistical significance at p<0.01, p<0.001, and p<0.0001, respectively).

#### 3.5.3. Pharmacological Treatment

For further validation of the functional capacity of our platform, we investigated the feasibility of this 3D in vitro model as a tool for drug screening applications. The EHTs were stimulated with two drug candidates with known effects, and their contractile activity was monitored before and after treatment. Norepinephrine (NE) is a common β-adrenergic agonist and can enhance the contraction force and beating frequency of cardiac cells, while Blebbistatin (Bleb) is a known myosin inhibitor that can adversely affect the contractility of CMs [14]. Tissue contractility was measured before and after drug treatment, and the EHTs showed the expected drug response (**Fig. 5F-H**, **Movie S11-12**). NE stimulation significantly increased contractile force, contractile stress, and beating rate, while the addition of Bleb resulted in complete inhibition of contraction (**Fig. 5I-K**). These results demonstrate the potency of our 3D model in predicting both the inotropic and chronotropic effects of the drugs in a physiologically relevant manner.

### 3.6. Cell-scale Measurements

Previous literature has highlighted the important role of cell mechanics in biological and biomedical systems, which significantly dictate cell fate and tissue function [68]. Different methods have been used to measure cell-generated forces, either at the single-cell level [36, 69] or the multi-cellular level [70, 71]. Despite several studies measuring the global stresses of cardiac tissue based on the deflection of structural supports (e.g., cantilevers), there is a lack of research on determining the local stresses of CMs in engineered heart tissues.

The eMSGs were synthesized by adding carboxylate-modified fluorescent polystyrene particles to the ultrasoft polyacrylamide composition. These bright particles facilitate imaging of the borders of the eMSGs without light scattering through the thick tissue (**Fig. 6A**). This ability is crucial for live imaging, tracking the sensors, determining the precise shape of the sensors, and further calculating stresses at different time points without inducing phototoxicity and photobleaching. Additionally, the cell-scale size of these ultrasoft mechanosensors and their suitable compliance facilitated the measurement of cell-generated stresses within microtissues. It allowed for the analysis of their spatial patterns, which we refer to as cell-scale stress measurements.

**Fig. 6.**
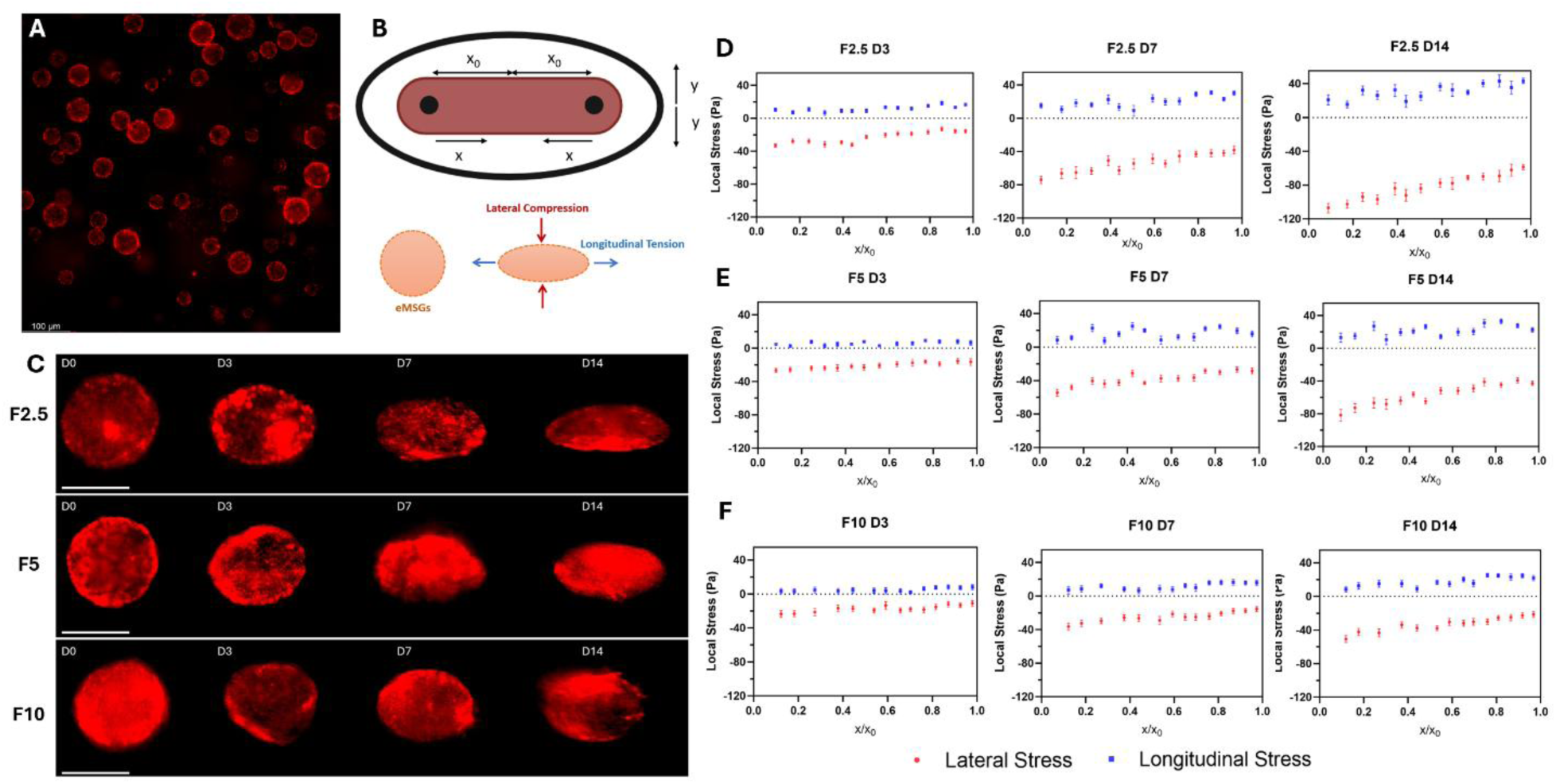
Stress measurements in 2D. (A) Synthesized eMSGs. (B) Schematic representation of the 2D system with longitudinal (x-axis) and lateral (y-axis) directions, as well as sensor deformation due to lateral compression and longitudinal tension. (C) Representative images of sensor deformations in F2.5, F5, and F10 microtissues at day 0, and after 3, 7, and 14 days. Scale bar = 50 μm. Longitudinal (blue) and lateral (red) internal stresses in F2.5 (D), F5 (E), and F10 (F) microtissues at different locations (x_0_ is half the interpillar distance).

#### 3.6.1. Stress measurements

The functional assessments of our EHTs demonstrated the significant potential of this platform as an in vitro model. However, our previous results only indicated global contractility at the tissue scale. We aimed to evaluate local contractility at the cellular scale and determine whether the same stress patterns could be observed across different matrix compositions and stiffness levels, based on our two measurement methods (pillar deflection and sensor deformation). eMSGs were encapsulated in the cell-hydrogel mixture and seeded in the chamber of HOCs. The concentration of the eMSG dispersion was adjusted to ensure enough traceable beads in each microtissue. The shape of each eMSG was monitored and recorded at four specific time points: immediately after tissue formation (Day 0), three days post-seeding (Day 3), one week of culture (Day 7), and two weeks of culture (Day 14). The reason for selecting these time intervals lies in tissue compaction and contraction trends. As explained previously, the cells remodeled the hydrogel into a compacted tissue strip after three days of culture, and the tissue started beating spontaneously. Tissue beating was observed up to Day 7, after which the tissue exhibited only passive compaction with no further contractile activity. Due to the complexity of our HOC system and the spatial 3D tissue compaction and contraction, specific logical assumptions were made to simplify the 3D model into a 2D Cartesian model. As shown in **Fig. 6B**, the direction of tissue elongation along the two pillars was defined as the x-axis, the orientation of lateral compaction as the y-axis, and the direction of tissue thickness as the z-axis. Based on our previous observations regarding eMSGs in spheroids, we assumed these sensors indicate elliptical deformation in 3D and exhibit y-z symmetry. This assumption significantly facilitated our strain-stress calculations, as our live imaging was performed with an inverted wide-field microscope, which depicted the shape of the sensors in the x-y orientation. Therefore, we could utilize our custom-built MATLAB code, modeling eMSGs as spherical bodies of a Neo-Hookean material in a homogeneous, nonlinear, and 2D axisymmetric finite element simulation. The evolution of sensor deformation was monitored in different ECM compositions (F2.5, F5, and F10 microtissues) over two weeks of culture. This deformation progression is depicted in **Fig. 6C** for representative eMSGs in these three microtissues. Each sensor became more compressed laterally and more elongated longitudinally, changing its shape from spherical to ellipsoidal. The elongated sensors aligned in the same direction as the cells, and the lateral compression followed the same orientation as tissue compaction. The sensor deformation results were further quantified for longitudinal stress (x-axis) and lateral stress (y-axis). Compressive and tensile stresses were indicated as negative and positive values, respectively. The internal stress calculations demonstrated more negative lateral stress and more positive longitudinal stress for each sensor over time (**Fig. 6D-F**). The change in lateral stresses was more significant due to progressive lateral compression over time (in the y-axis). Local heterogeneity in stresses within each tissue was observed (by changing x and y), as higher lateral stress was also observed in areas adjacent to the pillars compared to the center of the tissues, which is consistent with previous reports [20, 51, 72]. Increasing the stress values from day 3 to day 7 could be attributed to higher tissue contractile stresses, while the increase in stress values from day 7 to day 14 could be explained by the domination of CFs over CMs (similar to the mechanobiology of fibrosis progression [43]). Interestingly, the stress measurements revealed higher stress values at lower fibrin concentrations, which could also be visualized by greater sensor elongation in these microtissues. These results suggest that the sensor deformations and cell-generated stresses arose from the synergistic effect of higher CF-mediated tissue compaction and CM-generated contractile stresses. Previous studies have highlighted the role of lateral compaction in tissue stiffening and enhanced compaction forces in engineered 3D microtissues [73–75]. In particular, for cardiac tissues, it has been suggested that higher tissue compaction in a less rigid ECM overrides other factors influencing the trend of generated contractile stress, as we observed in our study [21, 41]. More importantly, the trends in the eMSG results closely matched our tissue functional results based on pillar deflection and calcium transients.

#### 3.6.2. Error Assessment

Following tissue fixation for IF purposes, the samples were imaged under a confocal microscope to observe the 3D structure of eMSGs and test our initial hypothesis regarding their ellipsoidal shape. We first confirmed that tissue fixation did not affect eMSG shape by comparing sensors before and after fixation (**Fig. S9**). Sensor’s strain values varied by less than 10%, and we therefore concluded that fixation did not affect the stress readouts. This result was expected, as eMSGs do not degrade during fixation or tissue culture. Fixation with paraformaldehyde typically causes minimal size and shape changes in the tissue by crosslinking its structure, which also preserves the bead’s shape. Hence, the z-axis strains could be estimated based on the spherical shape of eMSGs at day 0.

As indicated in **Fig. 7A** and **Movie S13**, the sensors did not show complete y-z symmetry, and partial flatness was detected on one side of them. Therefore, since the 3D structure of the deformed sensors was somewhat between ellipsoidal and pancake-like configurations, there was an error associated with our preliminary stress calculations (**Fig. 6**). This asymmetry correlates with the non-uniform dimensions of the seeding chamber, as the total length of the chamber (x-axis) is twice the width (y-axis), and the width is twice the thickness (z-axis). This led to heterogeneous spatial tissue compaction, which in turn influenced the corresponding irregularity observed in 3D sensor deformation.

**Fig. 7.**
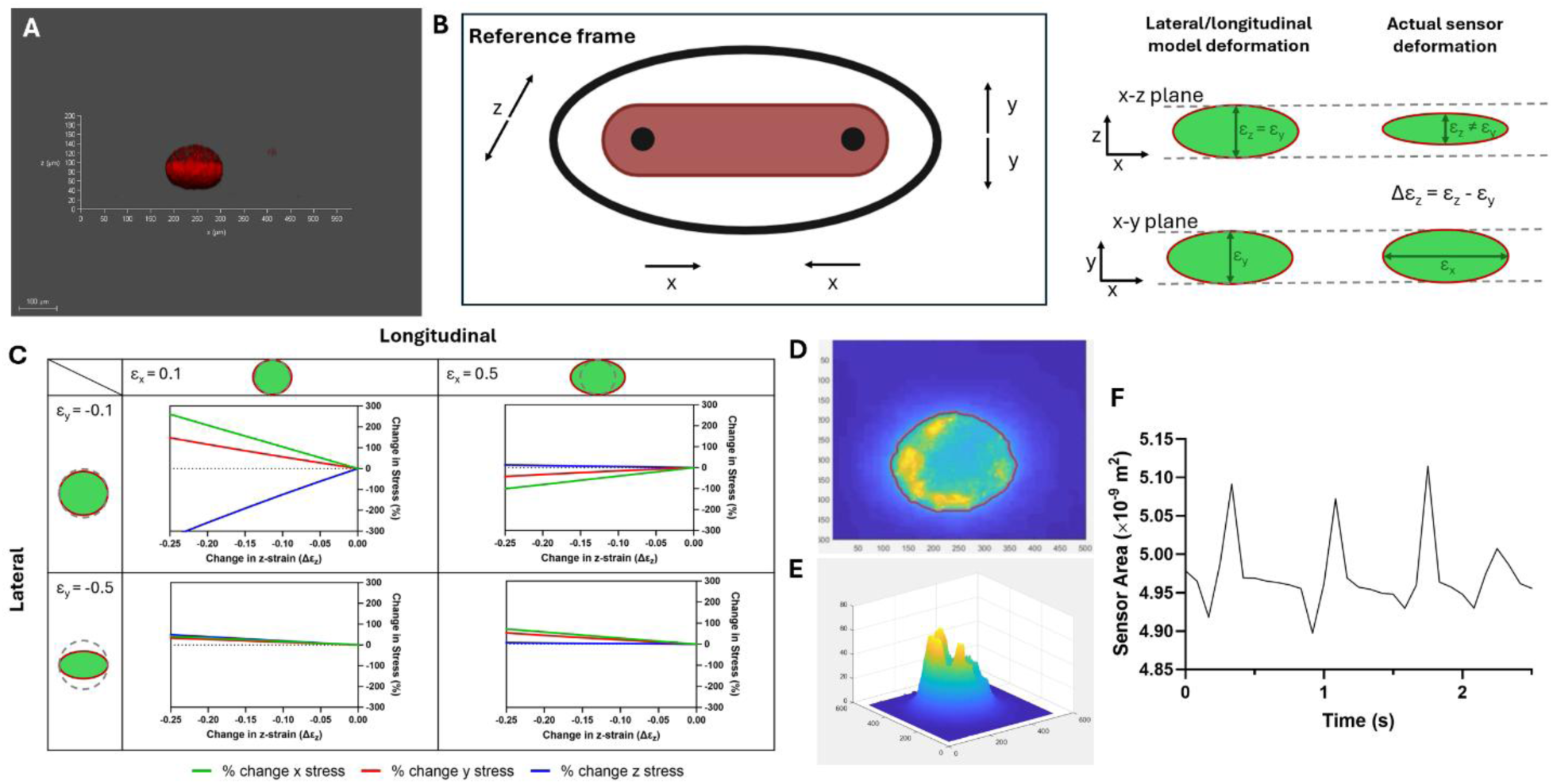
Error assessment of the 2D model and sensor deformation during tissue contraction. (A) Representative image of eMSGs in the z-x dimensions. (B) Schematic representation of the 3D model. (C) Error percentage in stresses based on strain values in the x and y directions. (D) Image processing of the sensors during a single beat showed successful detection of edges. (E) Optical flow analysis showed the area changes in the eMSGs. (F) Quantified amounts of the sensor area change during the systole/diastole cycle.

A third dimension was added to the sensor deformation modeling, indicating that the strains in the y- and z-axes were not identical (Δεz = εz – εy) (**Fig. 7B**). A finite element model was used to estimate the error that would arise in previous internal stress calculations (**Fig. S10**). Based on the measured strain values from 2D images, the approximate strain ranges were considered to be 0.1 to 0.5 in the longitudinal (εx) and −0.1 to −0.5 in the lateral (εy) directions, respectively. Z-strain was estimated to be ∼10-20% lower than the y-strain, representing a Δεz ∼ −0.2 to −0.1 (slightly more flat in the z-direction). Furthermore, an error of 20-40% was calculated for the previous model based on these strain values and the 3D model (**Fig. 7C** and **S11**). Smaller strains were associated with higher relative errors because smaller strains produce lower stress values, making stress deviations a more significant proportion of the original value and thereby increasing the relative error. Based on these calculations and considering the variabilities, we can conclude that the error margin is therefore reasonable and the trend of findings remains the same.

#### 3.6.3. Sensor Deformation during Contraction

Since the eMSGs are force-sensitive, we further investigated their shape alteration during tissue contraction using a fluorescent microscope. As shown in **Movie S14**, the sensor movement was clearly detectable, while changes in sensor shape were not distinguishable. The inability to detect these changes could be attributed to microscopic limitations, such as temporal resolution and capturing frame rates. To overcome these limitations, we analyzed the captured videos by applying a series of image processing techniques to enhance the subtle details and improve visualization and accuracy. First, pre-processing steps, including image stabilization, noise reduction, and contrast adjustment, were used to improve image clarity (**Movie S15**). Next, background subtraction and segmentation were applied to isolate regions of interest, followed by edge detection and thresholding to highlight subtle changes (**Fig. 7D** and **Movie S16**). Optical flow analysis and tracking algorithms were utilized to detect and quantify motion between frames (**Fig. 7E** and **Movie S17**). Finally, frame subtraction and time-lapse techniques amplified small variations over time, with intensity measurements and shape analysis providing quantitative insights into the observed changes and determining the sensor area changes during a single cell beating (systole/diastole cycle) (**Fig. 7F**). This detailed evaluation confirmed the sensor size changes during the contraction period, which could offer significant research potential in cardiac mechanobiology to correlate micro-scale local forces to global tissue contraction.

Our findings highlight the importance of multiscale mechanical analysis and raise key questions about the relationship between local and global contractile outputs in engineered cardiac tissues. Compared to conventional models, this dual-scale approach provides a more comprehensive contractile profile of EHTs and offers a unique opportunity to evaluate how individual cellular contractions contribute to overall tissue mechanics. We observed approximately an order-of-magnitude difference between the cell-scale stresses (measured by eMSGs) and the tissue-scale contraction (measured by pillar deflection). This difference arises because the eMSGs measure local, peak stresses generated at the cell-matrix interface, whereas the pillar deflection reflects the net global force transmitted through the tissue as a whole. The pillar force integrates contributions from all cells, with mechanical loads distributed across the tissue, resulting in higher net forces at the tissue level.

While the current study focuses on establishing and validating this dual-scale measurement approach, future work should explore how these two mechanical readouts correlate over time and across experimental conditions. The simultaneous measurement of both scales allows us to correlate local and global contractile behaviors, revealing insights into how cellular stresses contribute to overall tissue function. This is important because the global contraction measured by the pillar is not simply the sum of local cell contractions—it incorporates cell-cell interactions, matrix force transmission, and tissue-level coordination. By spatially mapping eMSG signals and modeling their contribution to global force output, this platform may ultimately allow derivation of global contractile stress without reliance on physical pillars, enabling more scalable and adaptable HOC systems.

## 4. Conclusion

In conclusion, our study presented an innovative cardiac microphysiological system by combining both flexible silicone pillars and eMSG sensor measurement techniques for simultaneous and continuous mapping of contractile stresses at both micro- and macro-scales. The model successfully supports tissue formation, compaction, and spontaneous beating of CMs, with robust functional and structural validation through immunofluorescence, calcium, and contractility assays. A pronounced effect of ECM composition and stiffness on cardiac tissue functionality was observed, with lower fibrin concentrations resulting in significantly higher beating frequency, contractile stress, and beating regularity. The model was further validated by demonstrating the chronotropic and inotropic effects of established drugs. The progression of incorporated eMSGs over two weeks of culture revealed that local cell-scale stresses followed the same trend as the global contractile stresses, enhanced by the synergistic effect of tissue compaction and tissue contraction. The platform’s capacity to assess local cell- and ECM-scale mechanics, combined with its high-throughput potential, offers a powerful tool for drug screening and mechanistic studies of cardiac function. This approach represents a significant advance in cardiac tissue engineering and pharmacological testing.

## Acknowledgment

Our work is supported by the Natural Sciences and Engineering Research Council of Canada (NSERC) (NSERC, RGPIN-2021-03960, DGECR-2021-00337 to HS, and RGPIN-2022-05165 to CM), Fonds de Recherche du Québec Santé (FRQS) (Chercheurs-boursiers J1 (313837)), Establishment of Young Investigators (324277), Montréal TransMedTech Institute (iTMT), CRCHU Sainte-Justine (CRCHUSJ), and the University of Montréal. A.M. gratefully acknowledges the FRQS Doctoral Scholarship, the Merit Scholarship from the Faculty of Medicine of the University of Montréal, and the Michel Bergeron Scholarship from the Department of Pharmacology and Physiology of the University of Montréal. C.-M. B. gratefully acknowledges support from the FRQNT and NSERC PGS-D doctoral awards. Access to COMSOL Multiphysics was provided by CMC microsystems.

## References

[1] B. Gu, K. Han, H. Cao, X. Huang, X. Li, M. Mao, H. Zhu, H. Cai, D. Li, J. He, Heart-on-a-chip systems with tissue-specific functionalities for physiological, pathological, and pharmacological studies, Materials Today Bio (2023) 100914.

[2] Y. Zhao, N. Rafatian, E.Y. Wang, Q. Wu, B.F. Lai, R.X. Lu, H. Savoji, M. Radisic, Towards chamber specific heart-on-a-chip for drug testing applications, Advanced drug delivery reviews 165 (2020) 60–76.

[3] C.A. Blair, B.L. Pruitt, Mechanobiology assays with applications in cardiomyocyte biology and cardiotoxicity, Advanced healthcare materials 9(8) (2020) 1901656.

[4] R. Portillo-Lara, A.R. Spencer, B.W. Walker, E.S. Sani, N. Annabi, Biomimetic cardiovascular platforms for in vitro disease modeling and therapeutic validation, Biomaterials 198 (2019) 78–94.

[5] H. Savoji, M.H. Mohammadi, N. Rafatian, M.K. Toroghi, E.Y. Wang, Y. Zhao, A. Korolj, S. Ahadian, M. Radisic, Cardiovascular disease models: a game changing paradigm in drug discovery and screening, Biomaterials 198 (2019) 3–26.

[6] M.L. McCain, H. Yuan, F.S. Pasqualini, P.H. Campbell, K.K. Parker, Matrix elasticity regulates the optimal cardiac myocyte shape for contractility, American Journal of Physiology-Heart and Circulatory Physiology 306(11) (2014) H1525–H1539.

[7] N. Sun, M. Yazawa, J. Liu, L. Han, V. Sanchez-Freire, O.J. Abilez, E.G. Navarrete, S. Hu, L. Wang, A. Lee, Patient-specific induced pluripotent stem cells as a model for familial dilated cardiomyopathy, Science translational medicine 4(130) (2012) 130ra47–130ra47.

[8] A.J. Ribeiro, O. Schwab, M.A. Mandegar, Y.-S. Ang, B.R. Conklin, D. Srivastava, B.L. Pruitt, Multi-imaging method to assay the contractile mechanical output of micropatterned human iPSC-derived cardiac myocytes, Circulation research 120(10) (2017) 1572–1583.

[9] N. Hu, T. Wang, Q. Wang, J. Zhou, L. Zou, K. Su, J. Wu, P. Wang, High-performance beating pattern function of human induced pluripotent stem cell-derived cardiomyocyte-based biosensors for hERG inhibition recognition, Biosensors and Bioelectronics 67 (2015) 146–153.

[10] J.H. Tsui, A. Leonard, N.D. Camp, J.T. Long, Z.Y. Nawas, R. Chavanachat, A.S. Smith, J.S. Choi, Z. Dong, E.H. Ahn, Tunable electroconductive decellularized extracellular matrix hydrogels for engineering human cardiac microphysiological systems, Biomaterials 272 (2021) 120764.

[11] M. Dong, N.-E. Oyunbaatar, P.P. Kanade, D.-S. Kim, D.-W. Lee, Real-time monitoring of changes in cardiac contractility using silicon cantilever arrays integrated with strain sensors, ACS sensors 6(10) (2021) 3556–3563.

[12] W. Dou, M. Malhi, Q. Zhao, L. Wang, Z. Huang, J. Law, N. Liu, C.A. Simmons, J.T. Maynes, Y. Sun, Microengineered platforms for characterizing the contractile function of in vitro cardiac models, Microsystems & Nanoengineering 8(1) (2022) 26.

[13] Y. Zhao, N. Rafatian, N.T. Feric, B.J. Cox, R. Aschar-Sobbi, E.Y. Wang, P. Aggarwal, B. Zhang, G. Conant, K. Ronaldson-Bouchard, A platform for generation of chamber-specific cardiac tissues and disease modeling, Cell 176(4) (2019) 913–927. e18.

[14] C. He, X. Wei, T. Liang, M. Liu, D. Jiang, L. Zhuang, P. Wang, Quantifying the compressive force of 3D cardiac tissues via calculating the volumetric deformation of built-in elastic gelatin microspheres, Advanced healthcare materials 10(16) (2021) 2001716.

[15] S.B. Bremner, K.S. Gaffney, N.J. Sniadecki, D.L. Mack, A change of heart: Human cardiac tissue engineering as a platform for drug development, Current Cardiology Reports 24(5) (2022) 473–486.

[16] J. Criscione, Z. Rezaei, C.M.H. Cantu, S. Murphy, S.R. Shin, D.-H. Kim, Heart-on-a-chip platforms and biosensor integration for disease modeling and phenotypic drug screening, Biosensors and Bioelectronics 220 (2023) 114840.

[17] K.W. Cho, W.H. Lee, B.-S. Kim, D.-H. Kim, Sensors in heart-on-a-chip: A review on recent progress, Talanta 219 (2020) 121269.

[18] C.-P. Heisenberg, Y. Bellaïche, Forces in tissue morphogenesis and patterning, Cell 153(5) (2013) 948–962.

[19] R. Riahi, S.A.M. Shaegh, M. Ghaderi, Y.S. Zhang, S.R. Shin, J. Aleman, S. Massa, D. Kim, M.R. Dokmeci, A. Khademhosseini, Automated microfluidic platform of bead-based electrochemical immunosensor integrated with bioreactor for continual monitoring of cell secreted biomarkers, Scientific reports 6(1) (2016) 24598.

[20] H. Wang, A.A. Svoronos, T. Boudou, M.S. Sakar, J.Y. Schell, J.R. Morgan, C.S. Chen, V.B. Shenoy, Necking and failure of constrained 3D microtissues induced by cellular tension, Proceedings of the National Academy of Sciences 110(52) (2013) 20923–20928.

[21] T. Boudou, W.R. Legant, A. Mu, M.A. Borochin, N. Thavandiran, M. Radisic, P.W. Zandstra, J.A. Epstein, K.B. Margulies, C.S. Chen, A microfabricated platform to measure and manipulate the mechanics of engineered cardiac microtissues, Tissue Engineering Part A 18(9-10) (2012) 910–919.

[22] M.S. Hutson, Y. Tokutake, M.-S. Chang, J.W. Bloor, S. Venakides, D.P. Kiehart, G.S. Edwards, Forces for morphogenesis investigated with laser microsurgery and quantitative modeling, Science 300(5616) (2003) 145–149.

[23] I.A. Morales, C.-M. Boghdady, B.E. Campbell, C. Moraes, Integrating mechanical sensor readouts into organ-on-a-chip platforms, Frontiers in Bioengineering and Biotechnology 10 (2022) 1060895.

[24] W. Lee, C.-M. Boghdady, V. Lelarge, R.L. Leask, L. McCaffrey, C. Moraes, Ultrasoft edge-labelled hydrogel sensors reveal internal tissue stress patterns in invasive engineered tumors, Biomaterials 296 (2023) 122073.

[25] W. Lee, N. Kalashnikov, S. Mok, R. Halaoui, E. Kuzmin, A.J. Putnam, S. Takayama, M. Park, L. McCaffrey, R. Zhao, Dispersible hydrogel force sensors reveal patterns of solid mechanical stress in multicellular spheroid cultures, Nature communications 10(1) (2019) 144.

[26] S. Mok, S. Al Habyan, C. Ledoux, W. Lee, K.N. MacDonald, L. McCaffrey, C. Moraes, Mapping cellular-scale internal mechanics in 3D tissues with thermally responsive hydrogel probes, Nature communications 11(1) (2020) 4757.

[27] A.A. Lucio, A. Mongera, E. Shelton, R. Chen, A.M. Doyle, O. Campàs, Spatiotemporal variation of endogenous cell-generated stresses within 3D multicellular spheroids, Scientific reports 7(1) (2017) 12022.

[28] E. Mohagheghian, J. Luo, J. Chen, G. Chaudhary, J. Chen, J. Sun, R.H. Ewoldt, N. Wang, Quantifying compressive forces between living cell layers and within tissues using elastic round microgels, Nature communications 9(1) (2018) 1878.

[29] M.E. Dolega, M. Delarue, F. Ingremeau, J. Prost, A. Delon, G. Cappello, Cell-like pressure sensors reveal increase of mechanical stress towards the core of multicellular spheroids under compression, Nature communications 8(1) (2017) 14056.

[30] N. Traeber, K. Uhlmann, S. Girardo, G. Kesavan, K. Wagner, J. Friedrichs, R. Goswami, K. Bai, M. Brand, C. Werner, Polyacrylamide bead sensors for in vivo quantification of cell-scale stress in zebrafish development, Scientific reports 9(1) (2019) 17031.

[31] A. Mousavi, A. Hedayatnia, P.P. van Vliet, D.R. Dartora, N. Wong, N. Rafatian, A.M. Nuyt, C. Moraes, A. Ajji, G. Andelfinger, Development of photocrosslinkable bioinks with improved electromechanical properties for 3D bioprinting of cardiac BioRings, Applied Materials Today 36 (2024) 102035.

[32] E. Pagan, E. Stefanek, A. Seyfoori, M. Razzaghi, B. Chehri, A. Mousavi, P. Arnaldi, Z. Ajji, D.R. Dartora, S.M.H. Dabiri, A handheld bioprinter for multi-material printing of complex constructs, Biofabrication 15(3) (2023) 035012.

[33] M.A. Tamargo, T.R. Nash, S. Fleischer, Y. Kim, O.F. Vila, K. Yeager, M. Summers, Y. Zhao, R. Lock, M. Chavez, milliPillar: a platform for the generation and real-time assessment of human engineered cardiac tissues, ACS biomaterials science & engineering 7(11) (2021) 5215–5229.

[34] J.H. Ahrens, S.G. Uzel, M. Skylar-Scott, M.M. Mata, A. Lu, K.T. Kroll, J.A. Lewis, Programming cellular alignment in engineered cardiac tissue via bioprinting anisotropic organ building blocks, Advanced Materials 34(26) (2022) 2200217.

[35] S. Finkel, S. Sweet, T. Locke, S. Smith, Z. Wang, C. Sandini, J. Imredy, Y. He, M. Durante, A. Lagrutta, FRESH™ 3D bioprinted cardiac tissue, a bioengineered platform for in vitro pharmacology, APL bioengineering 7(4) (2023).

[36] E.M. Strohm, N.I. Callaghan, Y. Ding, N. Latifi, N. Rafatian, S. Funakoshi, I. Fernandes, C.J. Reitz, M. Di Paola, A.O. Gramolini, Noninvasive Quantification of Contractile Dynamics in Cardiac Cells, Spheroids, and Organs-on-a-Chip Using High-Frequency Ultrasound, ACS nano 18(1) (2023) 314–327.

[37] Q. Wu, R. Xue, Y. Zhao, K. Ramsay, E.Y. Wang, H. Savoji, T. Veres, S.H. Cartmell, M. Radisic, Automated fabrication of a scalable heart-on-a-chip device by 3D printing of thermoplastic elastomer nanocomposite and hot embossing, Bioactive Materials 33 (2024) 46–60.

[38] O. Marques, Practical image and video processing using MATLAB, John Wiley & Sons 2011.

[39] M. Ząbczyk, R.A. Ariëns, A. Undas, Fibrin clot properties in cardiovascular disease: from basic mechanisms to clinical practice, Cardiovascular research 119(1) (2023) 94–111.

[40] A. Jafari, Z. Ajji, A. Mousavi, S. Naghieh, S.A. Bencherif, H. Savoji, Latest advances in 3D bioprinting of cardiac tissues, Advanced materials technologies 7(11) (2022) 2101636.

[41] N.J. Kaiser, R.J. Kant, A.J. Minor, K.L. Coulombe, Optimizing blended collagen-fibrin hydrogels for cardiac tissue engineering with human iPSC-derived cardiomyocytes, ACS biomaterials science & engineering 5(2) (2018) 887–899.

[42] Z. Wang, S.J. Lee, H.-J. Cheng, J.J. Yoo, A. Atala, 3D bioprinted functional and contractile cardiac tissue constructs, Acta biomaterialia 70 (2018) 48–56.

[43] E.Y. Wang, N. Rafatian, Y. Zhao, A. Lee, B.F.L. Lai, R.X. Lu, D. Jekic, L. Davenport Huyer, E.J. Knee-Walden, S. Bhattacharya, Biowire model of interstitial and focal cardiac fibrosis, ACS central science 5(7) (2019) 1146–1158.

[44] A. Mahdy, N.R. Webster, Perioperative systemic haemostatic agents, British journal of anaesthesia 93(6) (2004) 842–858.

[45] M. Afshar Bakooshli, E.S. Lippmann, B. Mulcahy, N. Iyer, C.T. Nguyen, K. Tung, B.A. Stewart, H. van den Dorpel, T. Fuehrmann, M. Shoichet, A 3D culture model of innervated human skeletal muscle enables studies of the adult neuromuscular junction, Elife 8 (2019) e44530.

[46] R.K. Christensen, C. von Halling Laier, A. Kiziltay, S. Wilson, N.B. Larsen, 3D printed hydrogel multiassay platforms for robust generation of engineered contractile tissues, Biomacromolecules 21(2) (2019) 356–365.

[47] M.J. Mondrinos, F. Alisafaei, A.Y. Yi, H. Ahmadzadeh, I. Lee, C. Blundell, J. Seo, M. Osborn, T.-J. Jeon, S.M. Kim, Surface-directed engineering of tissue anisotropy in microphysiological models of musculoskeletal tissue, Science advances 7(11) (2021) eabe9446.

[48] K.A. Jansen, R.G. Bacabac, I.K. Piechocka, G.H. Koenderink, Cells actively stiffen fibrin networks by generating contractile stress, Biophysical journal 105(10) (2013) 2240–2251.

[49] A. Patino-Guerrero, J. Veldhuizen, W. Zhu, R.Q. Migrino, M. Nikkhah, Three-dimensional scaffold-free microtissues engineered for cardiac repair, Journal of Materials Chemistry B 8(34) (2020) 7571–7590.

[50] K. Ronaldson-Bouchard, K. Yeager, D. Teles, T. Chen, S. Ma, L. Song, K. Morikawa, H.M. Wobma, A. Vasciaveo, E.C. Ruiz, Engineering of human cardiac muscle electromechanically matured to an adult-like phenotype, Nature protocols 14(10) (2019) 2781–2817.

[51] W.R. Legant, A. Pathak, M.T. Yang, V.S. Deshpande, R.M. McMeeking, C.S. Chen, Microfabricated tissue gauges to measure and manipulate forces from 3D microtissues, Proceedings of the National Academy of Sciences 106(25) (2009) 10097–10102.

[52] N.J. Hogrebe, J.W. Reinhardt, K.J. Gooch, Biomaterial microarchitecture: a potent regulator of individual cell behavior and multicellular organization, Journal of Biomedical Materials Research Part A 105(2) (2017) 640–661.

[53] V. Schwach, R. Passier, Native cardiac environment and its impact on engineering cardiac tissue, Biomaterials science 7(9) (2019) 3566–3580.

[54] M. Tiburcy, J.E. Hudson, P. Balfanz, S. Schlick, T. Meyer, M.-L. Chang Liao, E. Levent, F. Raad, S. Zeidler, E. Wingender, Defined engineered human myocardium with advanced maturation for applications in heart failure modeling and repair, Circulation 135(19) (2017) 1832–1847.

[55] A. Mousavi, S. Mashayekhan, N. Baheiraei, A. Pourjavadi, Biohybrid oxidized alginate/myocardial extracellular matrix injectable hydrogels with improved electromechanical properties for cardiac tissue engineering, International journal of biological macromolecules 180 (2021) 692–708.

[56] B.M. Baker, B. Trappmann, W.Y. Wang, M.S. Sakar, I.L. Kim, V.B. Shenoy, J.A. Burdick, C.S. Chen, Cell-mediated fibre recruitment drives extracellular matrix mechanosensing in engineered fibrillar microenvironments, Nature materials 14(12) (2015) 1262–1268.

[57] S. Munawar, I.C. Turnbull, Cardiac Tissue Engineering: Inclusion of Non-cardiomyocytes for Enhanced Features, Frontiers in Cell and Developmental Biology 9 (2021) 653127.

[58] M. Seguret, P. Davidson, S. Robben, C. Jouve, C. Pereira, Q. Lelong, L. Deshayes, C. Cerveau, M. Le Berre, R.S.R. Ribeiro, A versatile high-throughput assay based on 3D ring-shaped cardiac tissues generated from human induced pluripotent stem cell-derived cardiomyocytes, Elife 12 (2024) RP87739.

[59] A. Mousavi, S. Vahdat, N. Baheiraei, M. Razavi, M.H. Norahan, H. Baharvand, Multifunctional conductive biomaterials as promising platforms for cardiac tissue engineering, ACS biomaterials science & engineering 7(1) (2020) 55–82.

[60] A. Mousavi, E. Stefanek, A. Jafari, Z. Ajji, S. Naghieh, M. Akbari, H. Savoji, Tissue-engineered heart chambers as a platform technology for drug discovery and disease modeling, Biomaterials Advances 138 (2022) 212916.

[61] F. Zhang, H. Cheng, K. Qu, X. Qian, Y. Lin, Y. Zhang, S. Qian, N. Huang, C. Cui, M. Chen, Continuous contractile force and electrical signal recordings of 3D cardiac tissue utilizing conductive hydrogel pillars on a chip, Materials Today Bio 20 (2023) 100626.

[62] N. Thavandiran, C. Hale, P. Blit, M.L. Sandberg, M.E. McElvain, M. Gagliardi, B. Sun, A. Witty, G. Graham, V.T. Do, Functional arrays of human pluripotent stem cell-derived cardiac microtissues, Scientific reports 10(1) (2020) 6919.

[63] M.E. Afshar, H.Y. Abraha, M.A. Bakooshli, S. Davoudi, N. Thavandiran, K. Tung, H. Ahn, H.J. Ginsberg, P.W. Zandstra, P.M. Gilbert, A 96-well culture platform enables longitudinal analyses of engineered human skeletal muscle microtissue strength, Scientific reports 10(1) (2020) 6918.

[64] S.R. Shin, C. Zihlmann, M. Akbari, P. Assawes, L. Cheung, K. Zhang, V. Manoharan, Y.S. Zhang, M. Yüksekkaya, K.t. Wan, Reduced graphene oxide-gelMA hybrid hydrogels as scaffolds for cardiac tissue engineering, Small 12(27) (2016) 3677–3689.

[65] X.-P. Li, K.-Y. Qu, B. Zhou, F. Zhang, Y.-Y. Wang, O.D. Abodunrin, Z. Zhu, N.-P. Huang, Electrical stimulation of neonatal rat cardiomyocytes using conductive polydopamine-reduced graphene oxide-hybrid hydrogels for constructing cardiac microtissues, Colloids and Surfaces B: Biointerfaces 205 (2021) 111844.

[66] J.A. Schaefer, R.T. Tranquillo, Tissue contraction force microscopy for optimization of engineered cardiac tissue, Tissue Engineering Part C: Methods 22(1) (2016) 76–83.

[67] I. Mannhardt, K. Breckwoldt, D. Letuffe-Brenière, S. Schaaf, H. Schulz, C. Neuber, A. Benzin, T. Werner, A. Eder, T. Schulze, Human engineered heart tissue: analysis of contractile force, Stem cell reports 7(1) (2016) 29–42.

[68] A.M. Obenaus, M.Y. Mollica, N.J. Sniadecki, (De) form and function: measuring cellular forces with deformable materials and deformable structures, Advanced healthcare materials 9(8) (2020) 1901454.

[69] K. Sheets, J. Wang, W. Zhao, R. Kapania, A.S. Nain, Nanonet force microscopy for measuring cell forces, Biophysical Journal 111(1) (2016) 197–207.

[70] Z. Ma, N. Huebsch, S. Koo, M.A. Mandegar, B. Siemons, S. Boggess, B.R. Conklin, C.P. Grigoropoulos, K.E. Healy, Contractile deficits in engineered cardiac microtissues as a result of MYBPC3 deficiency and mechanical overload, Nature biomedical engineering 2(12) (2018) 955–967.

[71] C. Wang, S. Koo, M. Park, Z. Vangelatos, P. Hoang, B.R. Conklin, C.P. Grigoropoulos, K.E. Healy, Z. Ma, Cardiac microtissues: Maladaptive contractility of 3D human cardiac microtissues to mechanical nonuniformity (Adv. Healthcare mater. 8/2020), Advanced Healthcare Materials 9(8) (2020) 2070024.

[72] O.J. Abilez, E. Tzatzalos, H. Yang, M.-T. Zhao, G. Jung, A.M. Zöllner, M. Tiburcy, J. Riegler, E. Matsa, P. Shukla, Passive stretch induces structural and functional maturation of engineered heart muscle as predicted by computational modeling, Stem cells 36(2) (2018) 265–277.

[73] R. Zhao, C.S. Chen, D.H. Reich, Force-driven evolution of mesoscale structure in engineered 3D microtissues and the modulation of tissue stiffening, Biomaterials 35(19) (2014) 5056–5064.

[74] J. Jilberto, S.J. DePalma, J. Lo, H. Kobeissi, L. Quach, E. Lejeune, B.M. Baker, D. Nordsletten, A data-driven computational model for engineered cardiac microtissues, Acta biomaterialia 172 (2023) 123–134.

[75] J. Alvarado, M. Sheinman, A. Sharma, F.C. MacKintosh, G.H. Koenderink, Force percolation of contractile active gels, Soft matter 13(34) (2017) 5624–5644.

