## Supporting Information for "Heart-On-a-Chip with Integrated Ultrasoft Mechanosensors for Continuous Measurement of Cell- and Tissue-scale Contractile Stresses"

*** Corresponding Author**

**Supplementary Figures**

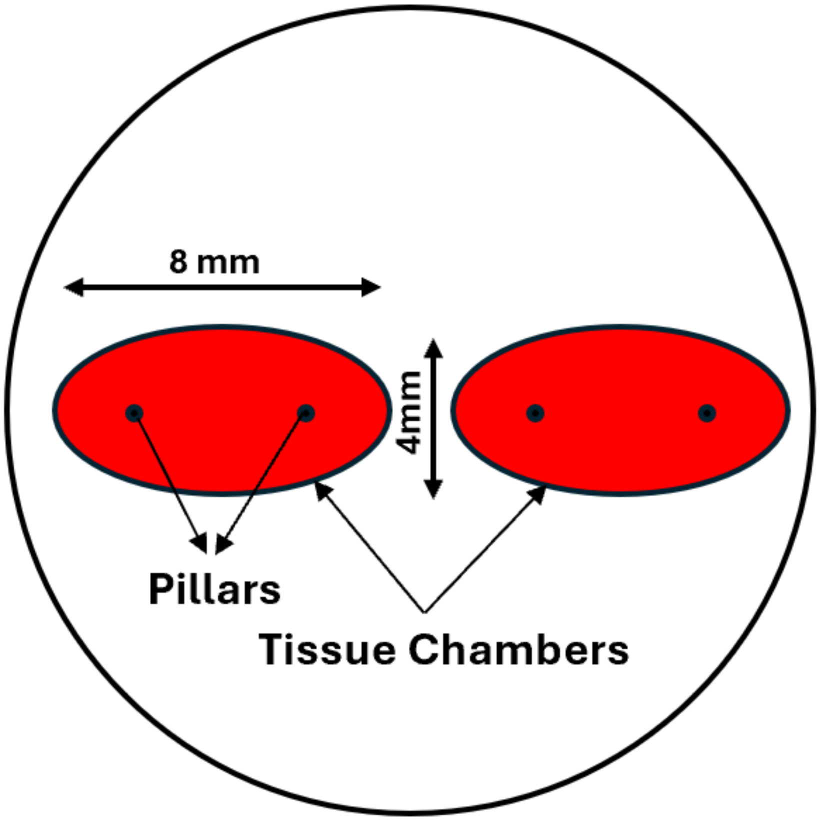


**Fig. S1.** Schematics of chip design and the dimensions.

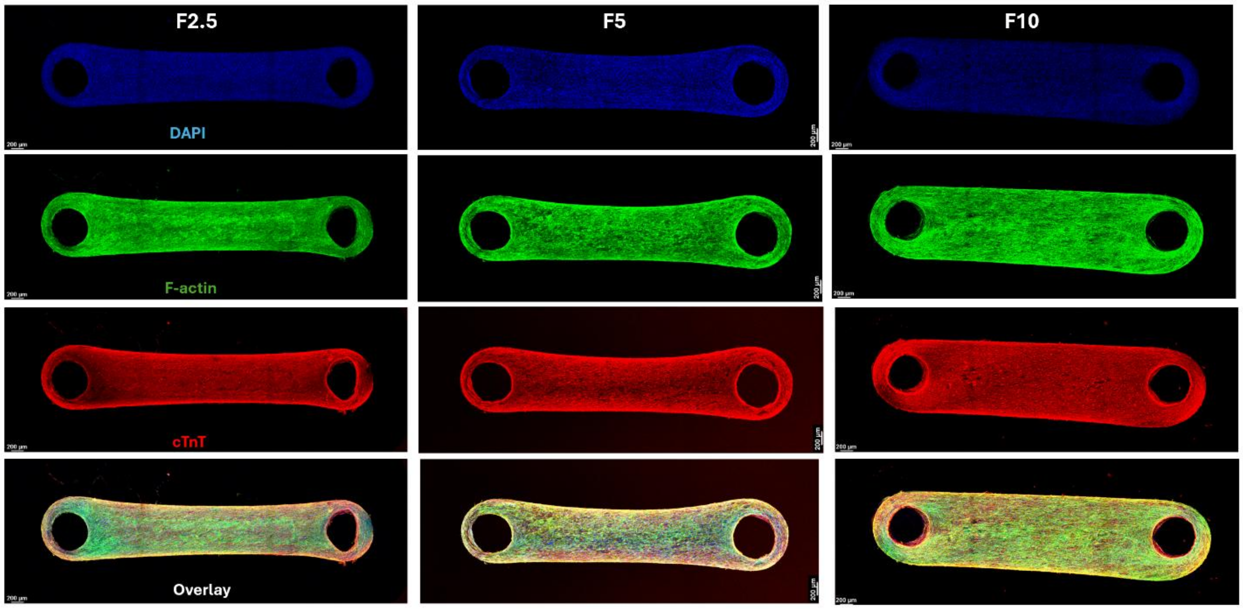


**Fig. S2.** Different channels of confocal imaging for the whole mount IF staining of F2.5, F5, and F10 cardiac microtissues after 7 days of culture (blue channel: DAPI, red channel: cTnT, green channel: phalloidin) and overlay images. Scale bar = 200 µm.

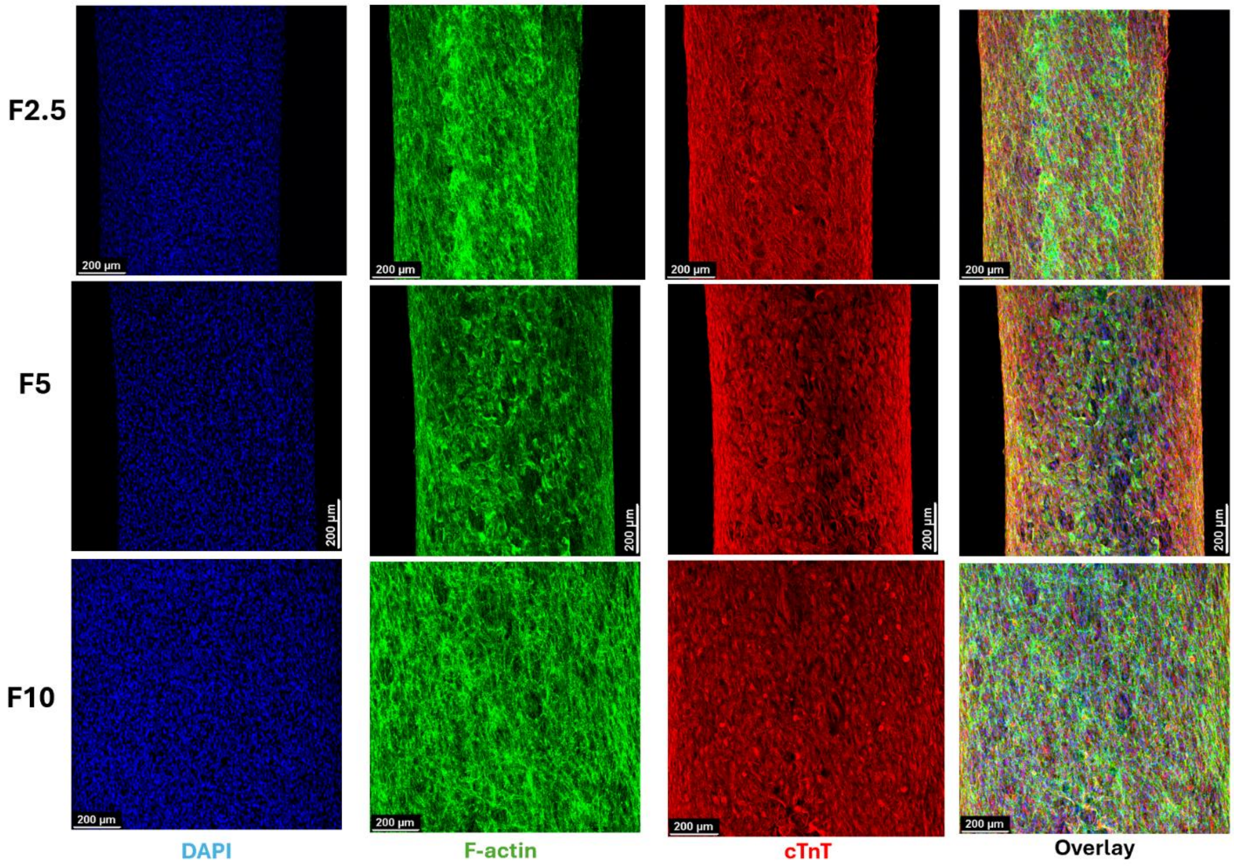


**Fig. S3.** Different channels of confocal imaging for the IF staining of F2.5, F5, and F10 cardiac microtissues (10x objective) after 7 days of culture (blue channel: DAPI, red channel: cTnT, green channel: phalloidin) and overlay images. Scale bar = 200 µm.

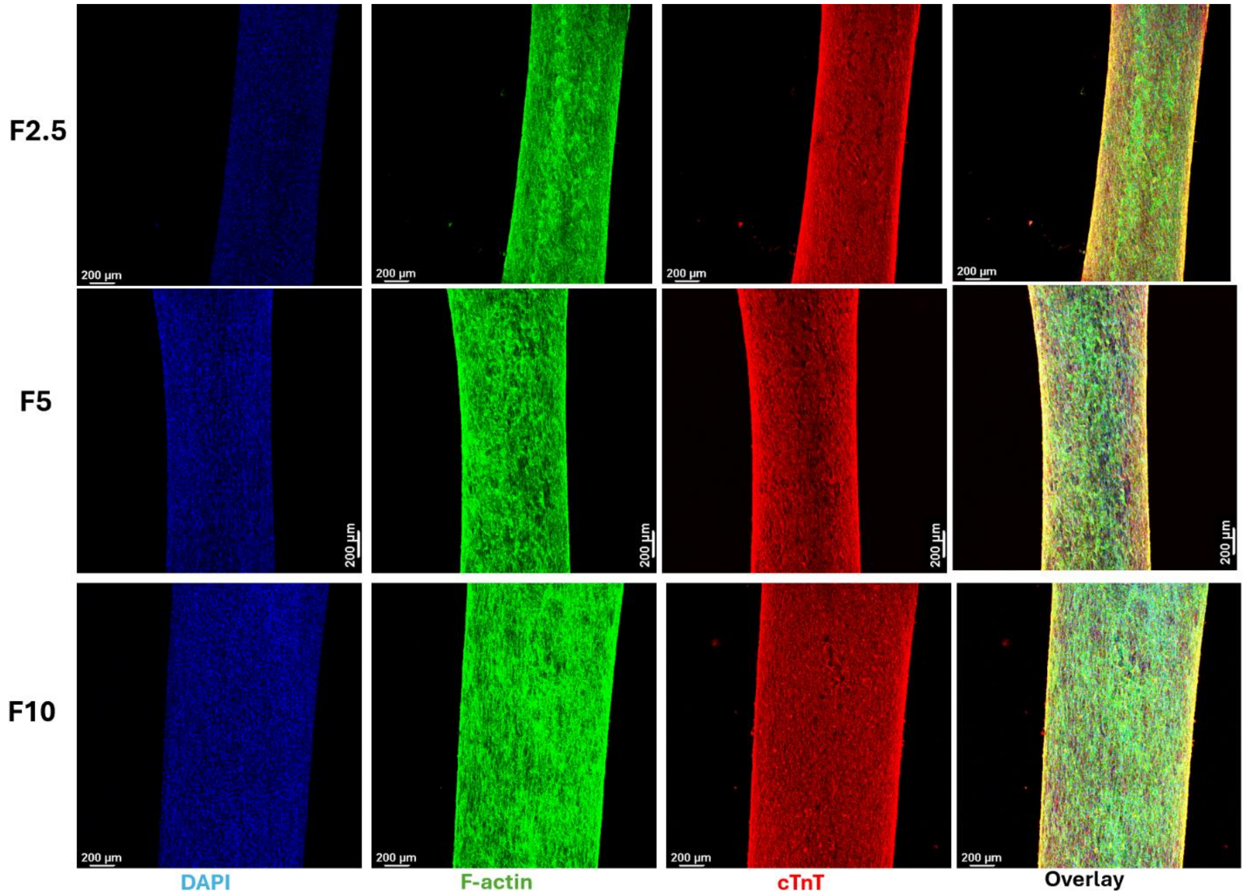


**Fig. S4.** Different channels of confocal imaging for the IF staining of F2.5, F5, and F10 cardiac microtissues (5x objective) after 7 days of culture (blue channel: DAPI, red channel: cTnT, green channel: phalloidin) and overlay images. Scale bar = 200 μm.

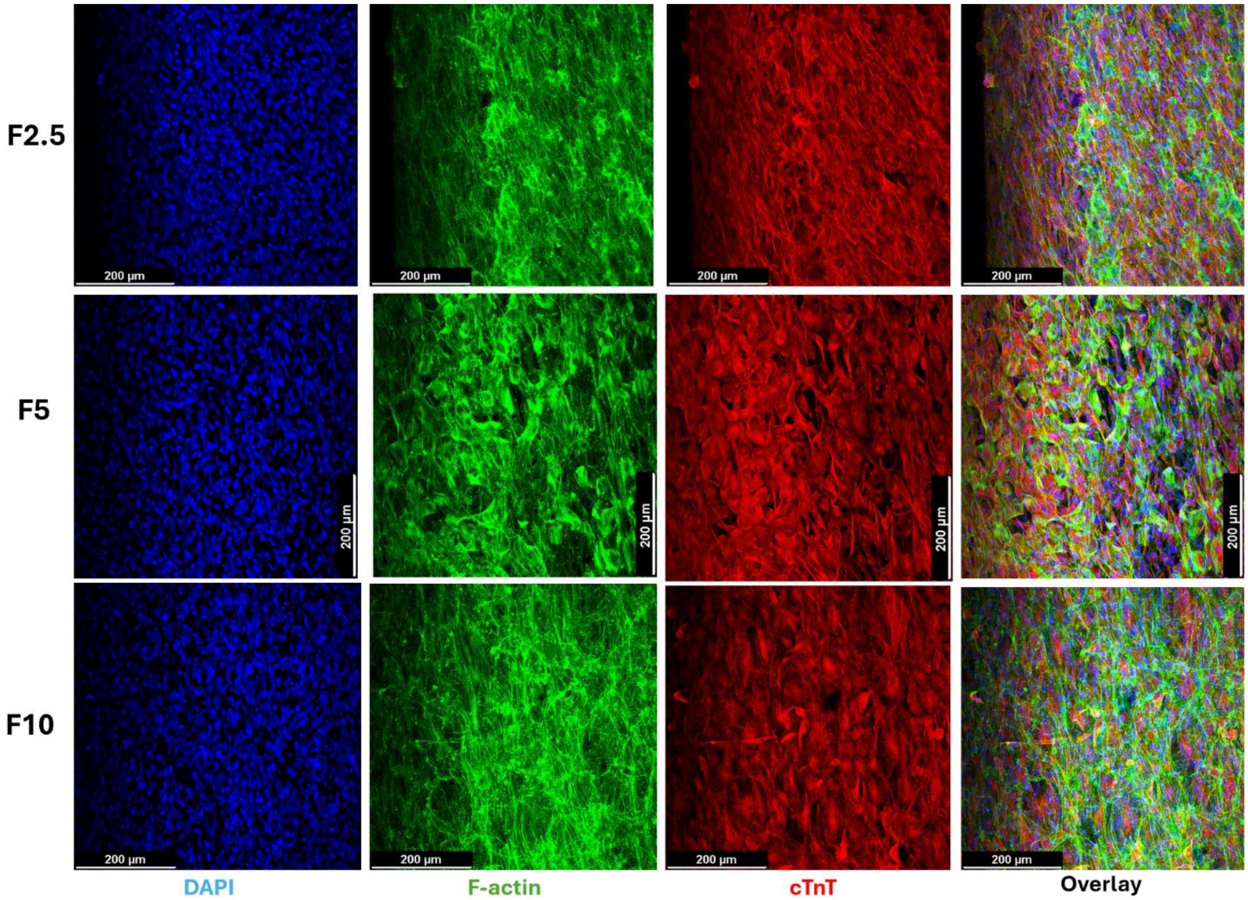


**Fig. S5.** Different channels of confocal imaging for the IF staining of F2.5, F5, and F10 cardiac microtissues (20x objective) after 7 days of culture (blue channel: DAPI, red channel: cTnT, green channel: phalloidin) and overlay images. Scale bar = 200 µm.

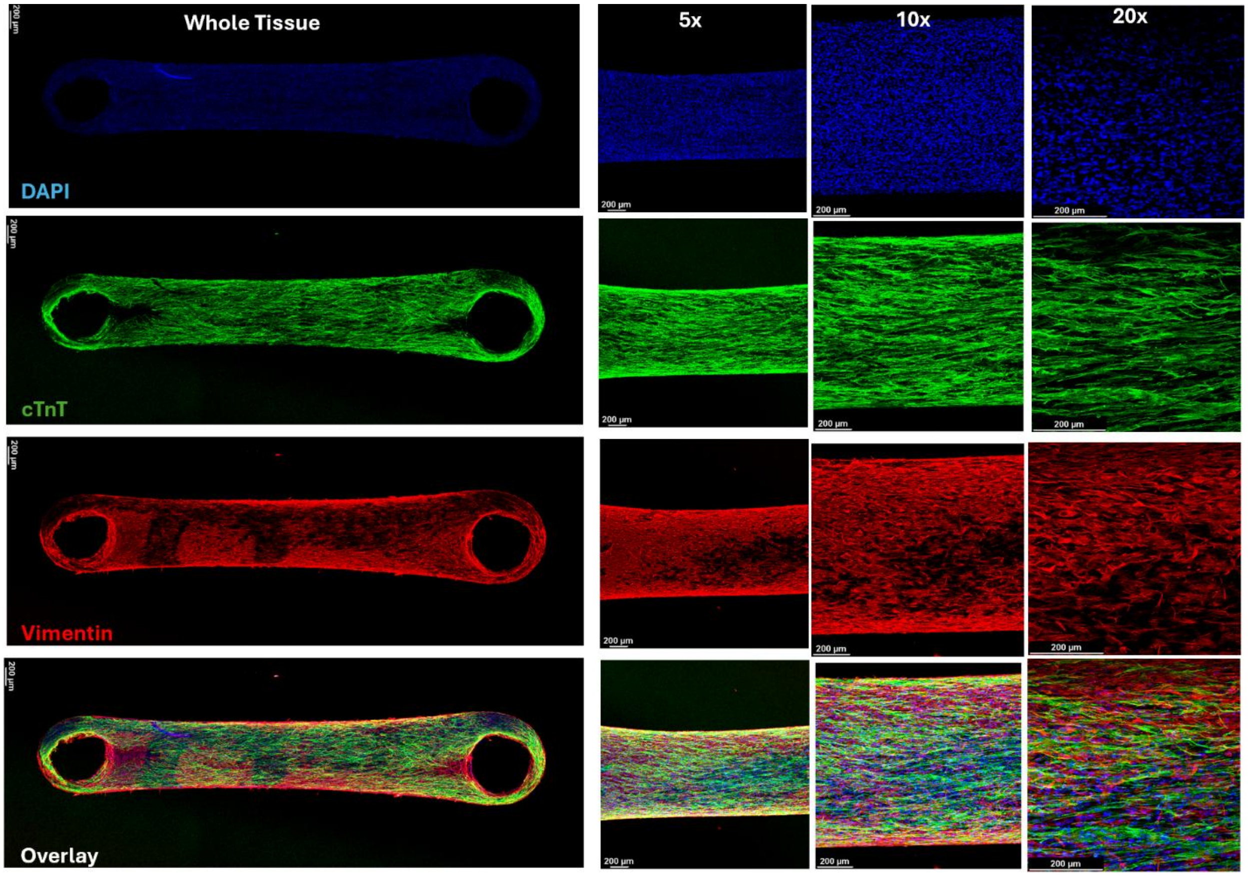


**Fig. S6.** Different channels of confocal imaging for the whole mount IF staining of cardiac microtissues for CMs vs non-CMs as well as different resolutions (5x, 10x, and 20x) after 7 days of culture (blue channel: DAPI, red channel: vimentin, green channel: cTnT) and overlay images. Scale bar = 200 µm.

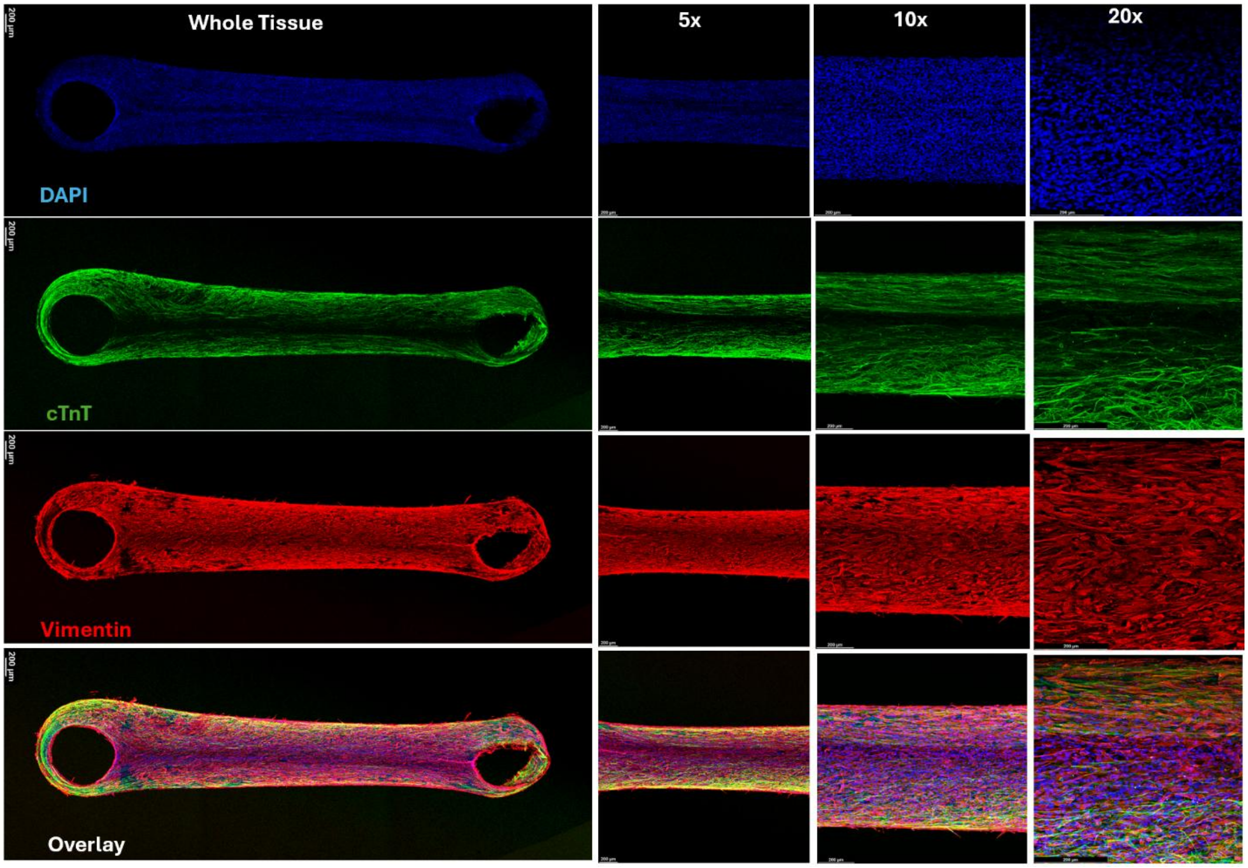


**Fig. S7.** Different channels of confocal imaging for the whole mount IF staining of cardiac microtissues for CMs vs non-CMs as well as different resolutions (5x, 10x, and 20x) after 14 days of culture (blue channel: DAPI, red channel: vimentin, green channel: cTnT) and overlay images. Scale bar = 200 µm.

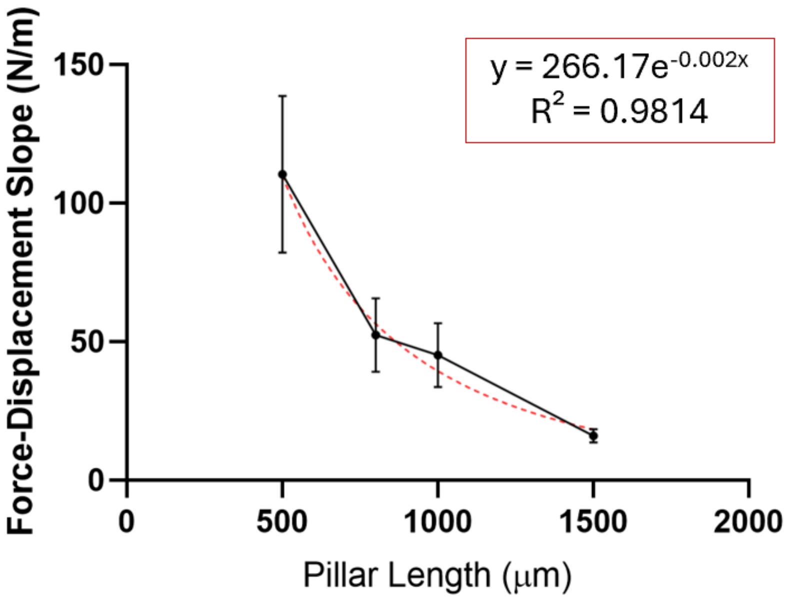


**Fig. S8.** Relationship between slop of force-displacement curve and pillar length (in the testing range of 0.5 to 1.5 mm).

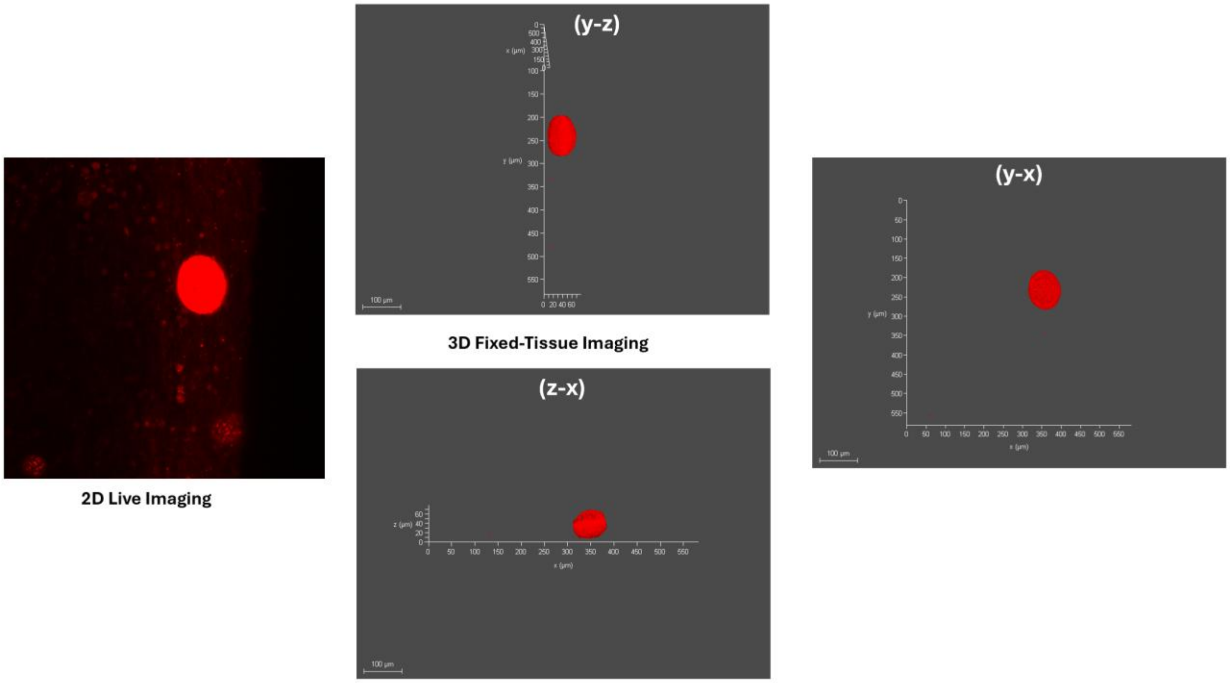


**Fig. S9.** Representative images of eMSGs before fixation (2D live imaging) and after fixation (3D confocal imaging) in y-z, z-x, and y-x orientations. Scale bar=100μm.

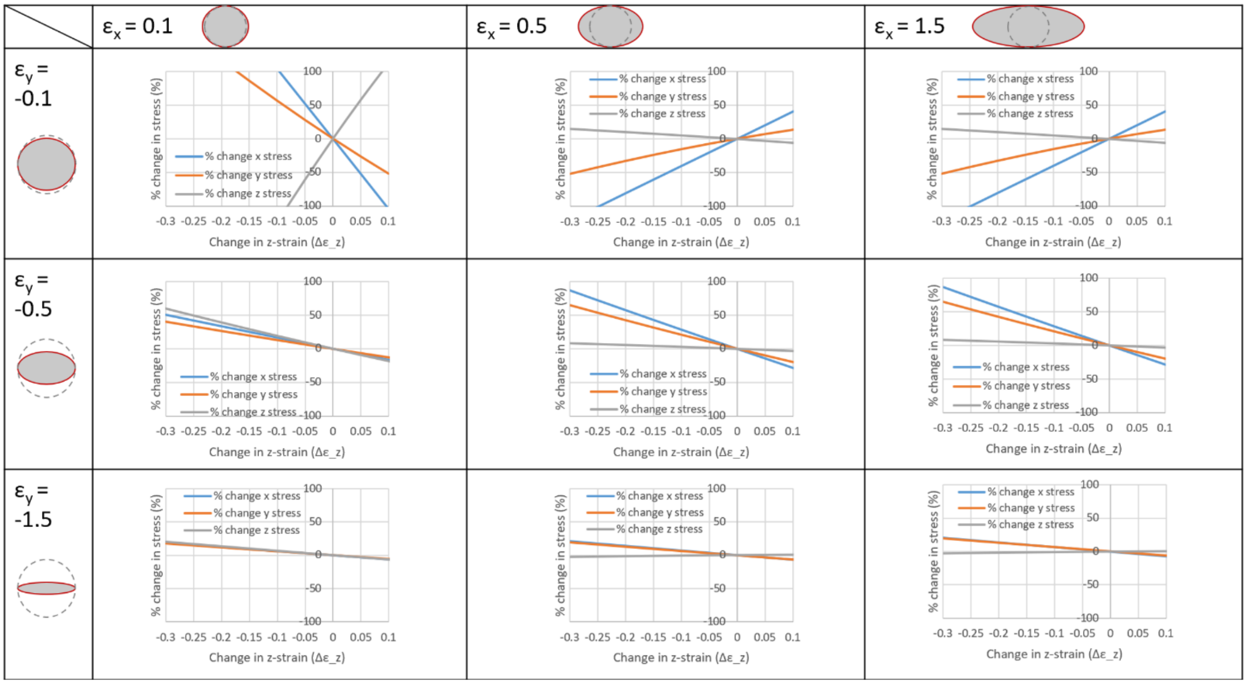


**Fig. S10.** COMSOL 3D modeling of the system with different strains in x and y orientations to estimate the error percentage of 2D stress measurements.

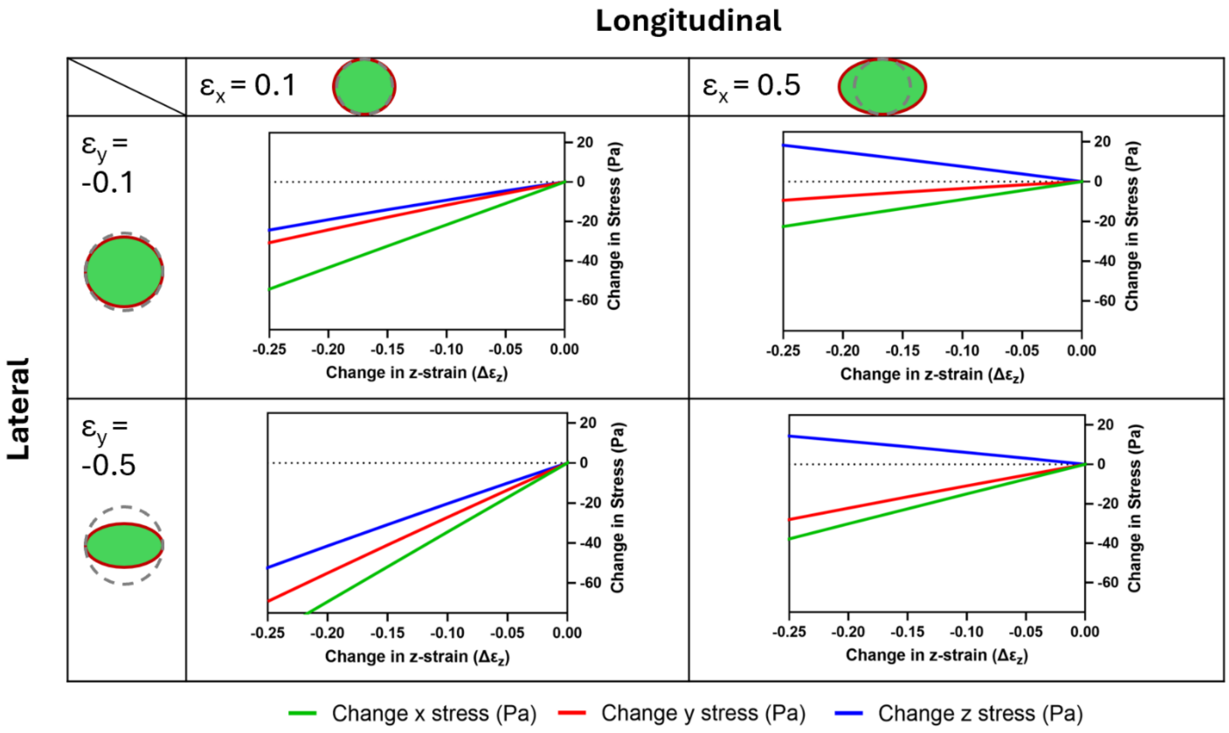


**Fig. S11.** COMSOL 3D modeling of the system to estimate the actual change in stresses (Pa) based on strain values in the x and y directions.

**Supplementary Movies**

**Movie S1:** Microsquisher analysis of pillars at 500 μm height from the base.

**Movie S2:** Microsquisher analysis of pillars at 800 μm height from the base.

**Movie S3:** Microsquisher analysis of pillars at 1000 μm height from the base.

**Movie S4:** Microsquisher analysis of pillars at 1500 μm height from the base.

**Movie S5:** Spontaneous beating of F2.5 cardiac microtissues on day 7 of culture.

**Movie S6:** Spontaneous beating of F5 cardiac microtissues on day 7 of culture.

**Movie S7:** Spontaneous beating of F10 cardiac microtissues on day 7 of culture.

**Movie S8:** Calcium transients of F5 cardiac microtissues on day 7 of culture.

**Movie S9:** Calcium transients of F10 cardiac microtissues on day 7 of culture.

**Movie S10:** Calcium transients of F5 cardiac microtissues on day 14 of culture.

**Movie S11:** Spontaneous beating of EHTs before the drug treatment (CTRL group).

**Movie S12:** Spontaneous beating of EHTs after the treatment with norepinephrine (NE group).

**Movie S13:** 3D view of a representative sensor by confocal imaging after tissue fixation.

**Movie S14:** A representative of video showing sensor movement during tissue contraction.

**Movie S15:** Image processing of Movie S14 to identify sensor deformation during contraction, including image stabilization, noise reduction, and contrast adjustment

**Movie S16:** Image processing of Movie S14 to identify sensor deformation during contraction, including background subtraction and segmentation, followed by edge detection and thresholding

**Movie S16:** Image processing of Movie S14 to identify sensor deformation during contraction, including background subtraction and segmentation, followed by edge detection and thresholding

**Movie S17:** Image processing of Movie S14 to identify sensor deformation during contraction, including optical flow analysis and the application of tracking algorithms to quantify motion between frames.
