## Supplementary figures and images for "Heart-On-a-Chip with Integrated Ultrasoft Mechanosensors for Continuous Measurement of Cell- and Tissue-scale Contractile Stresses"

### Movie S15

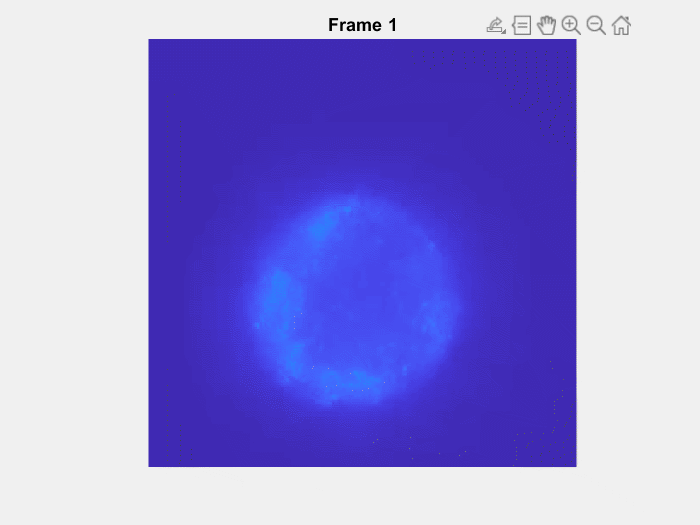

### Movie S16

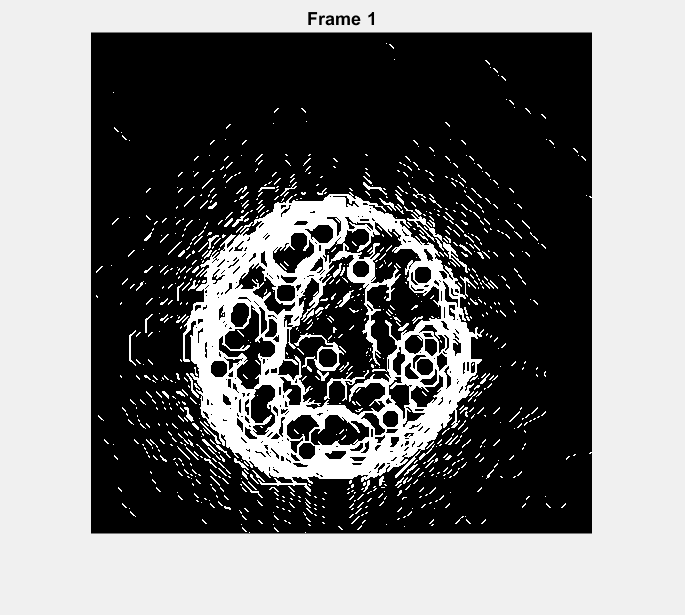

### Movie S17

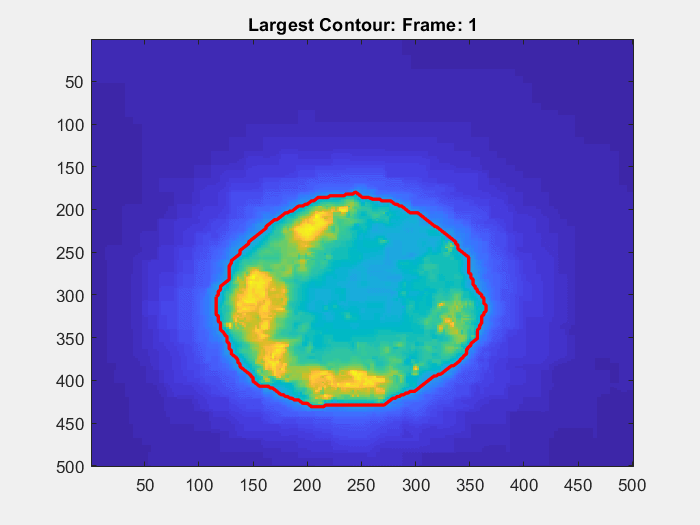
